# GPC3-Unc5D complex structure and role in cell migration

**DOI:** 10.1101/2022.07.21.500812

**Authors:** O Akkermans, C Delloye-Bourgeois, C Peregrina, M Carrasquero-Ordaz, M Kokolaki, M Berbeira-Santana, M Chavent, F Reynaud, Ritu Raj, J Agirre, M Aksu, E White, E Lowe, D Ben Amar, S Zaballa, J Huo, P.T.N. McCubbin, D Comoletti, R Owens, C.V. Robinson, V Castellani, D del Toro, E Seiradake

## Abstract

Neural migration is a critical step during brain development that requires the interactions of cell-surface guidance receptors. Cancer cells often hijack these mechanisms to disseminate. Here we reveal crystal structures of Uncoordinated-5 receptor D (Unc5D) in complex with morphogen receptor glypican-3 (GPC3), forming an octameric glycoprotein complex. In the complex, four Unc5D molecules pack into an antiparallel bundle, flanked by four GPC3 molecules. Central glycan-glycan interactions are formed by N-linked glycans emanating from GPC3 (N241 in human) and C-mannosylated tryptophans of the Unc5D thrombospondin-like domains. MD simulations, mass-spectrometry and structure-based mutants validate the crystallographic data. Anti-GPC3 nanobodies enhance or weaken Unc5-GPC3 binding. Using these tools *in vivo*, we show that Unc5/GPC3 guide migrating pyramidal neurons in the mouse cortex, and cancer cells in an embryonic xenograft neuroblastoma model. The results demonstrate a conserved structural mechanism of cell-guidance, with the potential for wide-ranging biomedical implications in development and cancer biology.

## Introduction

Context-dependent signalling networks formed by adhesion molecules, morphogens, guidance cues, and their receptors, direct cellular navigation, differentiation, and synapse formation. Guidance receptors of the Uncoordinated-5 family (Unc5A-D) have emerged as key players in navigating cells and axons (Hong et al., 1999; Leung-Hagesteijn et al., 1992), where they are known for their role in triggering repulsion in response to different ligands such as Fibronectin Leucine-rich Repeat Transmembrane proteins (FLRT1-3) (Seiradake et al., 2014; Yamagishi et al., 2011) and Netrins (Hong et al., 1999). We and others have previously shown that Unc5D guides the radial migration of excitatory neurons during cortical development, a key process that is required for the formation of functionally distinct cortical layers (Miyoshi and Fishell, 2012; Seiradake et al., 2014; Yamagishi et al., 2011). Pyramidal neurons born from the germinal layers of the mouse cortex, are initially multipolar while moving radially from the subventricular zone (SVZ) and through the intermediate zone (IZ). These cells transition to a bipolar morphology in the upper part of the IZ, attaching to the fibers of apical progenitor (AP) cells to enter the cortical plate (CP) and settle in their appropriate layer (Tabata and Nakajima, 2003). Relatively little is known about the molecular mechanisms that underpin the switch from slow multipolar to fast bipolar migration displayed by these neurons. Unc5D is one of the few molecular components that has been identified to regulate this process. Unc5D expression increases during the migration of multipolar neurons and decreases after these cells transition to a bipolar morphology (Miyoshi and Fishell, 2012). Altering Unc5D expression levels disrupts multipolar to bipolar transition, delays cortical migration and affects the layering position of cortical neurons (Miyoshi and Fishell, 2012). Unc5D and DCC are the only Netrin receptors expressed in the IZ. However, Netrin expression is low in this region during cortex development, suggesting a dependence on other ligands. Indeed, premature radial migration of Unc5D-positive neurons is prevented by a gradient of the repulsive ligand FLRT2, which is shed from cells in the CP towards the IZ (Yamagishi et al., 2011). Unc5 receptors have a conserved domain architecture, consisting of two extracellular immunoglobulin domains (Ig1-2), two extracellular thrombospondin-like domains (TSP1-2), a single transmembrane helix, and a C-terminal intracellular supramodule, which contains a ZO-1/Unc5 (ZU5), a Unc5/PIDD/Ankyrin (UPA), and a death domain (DD) (Fig. 1A). We previously solved the ectodomain structure of human Unc5A isoform 1 which lacks TSP1 (Seiradake et al., 2014) and rat Unc5D Ig1-Ig2-TSP1 in a super-complex with mouse FLRT2 and Latrophilin 3 (Jackson et al., 2016). Both structures revealed an elongated Unc5 molecule with a linear arrangement of Ig and TSP domains, with a FLRT binding site at the N-terminal Ig1 domain. The crystal structure of the Unc5B cytoplasmic domains ZU5-UPA-DD is also known and revealed a closed configuration that is thought to be in an autoinhibitory state (Wang et al., 2009).

**Figure 1.**
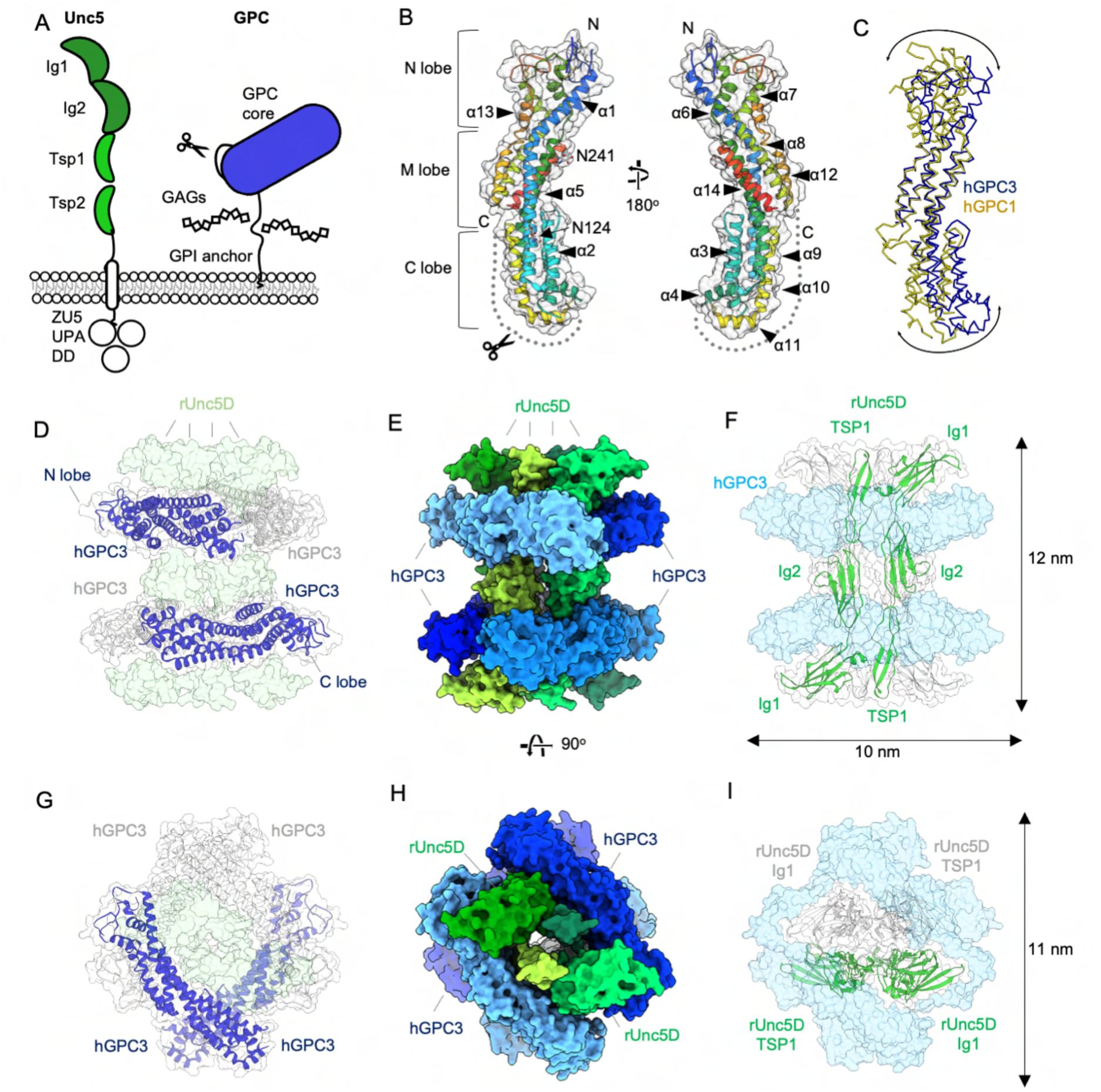
GPC3^core^ and Unc5^IgIgTSP^ crystal structures reveal an octameric complex. **(A)** Domain architecture of Unc5 and GPC3. A furin-type cleavage site is indicated by a scissors symbol. **(B)** Crystal structure of hGPC3^core^, using the nomenclature presented in (Kim et al. PNAS 2011). The structure is coloured according to the rainbow (blue = N-terminus, red= C-terminus). **(C)** Superposition of hGPC1, yellow (Awad et al. 2015), and hGPC3 core, blue. **(D-I)** Views of the hGPC3^core^-rUnc5^IgIgTSP^ complex in two orientations, indicating hGPC3^core^ as ribbons in dark blue and grey (D, G), an overview of the complex in solid surface view with GPC3 in shades of blue and Unc5 in shades of green (E, H), and views in which the Unc5 chains are indicated in green and grey ribbons (F, I).

Here we show that Unc5 receptors directly and functionally interact with the morphogen receptor glypican-3 (GPC3). Glypicans (GPC1-6 in mammals) are subdivided into two subfamilies based on sequence identity: GPC1/2/4/6 form one group with 38-64% sequence identity; GPC3/5 form the second group with ∼42% sequence identity (in humans). All glypicans share a similar molecular architecture, including a structured N-terminal N-glycosylated extracellular domain, the α-helical core, followed by a C-terminal linker region of ∼80 amino acids (Kim et al., 2011) (Fig 1A). Crystal structures of the core domain of human GPC1 (Awad et al., 2015; Svensson et al., 2012) and the fly ortholog Dally-Like-Protein (DLP) (Kim et al., 2011; McGough et al., 2020) have revealed a conserved α-helical cylindrical architecture comprising an N-terminal lobe (N lobe) that contains 6 conserved disulfide bonds, a central M lobe made up of 5 main helices (α1, α5, α8, α12, α14), and a C-terminal lobe (C lobe) that contains a furin-like convertase cleavage site (RXXR) between α11 and α12 (de Cat et al., 2003). Cleavage of this site results in the formation of two fragments that remain covalently attached through disulfide bonds (de Cat et al., 2003). The second conserved region, the C-terminal linker, carries a glycosylphosphatidylinositol (GPI) anchor that tethers the protein to the cell surface, and is an attachment site for heparan sulphate (HS) glycans (David et al., 1990; Watanabe et al., 1995) (Fig 1A). HS glycans are sufficient for, or contribute to, the binding of many reported GPC3 interaction partners (Wang et al., 2020), which include Wnts, the Wnt-receptor Frizzled and Hedgehog (Capurro et al., 2014, 2005, 2008). Structures to show how these proteins interact with GPC3 have not been reported.

Mutations in GPC3 cause Simpson-Golabi-Behmel overgrowth syndrome, a genetic disorder that presents with visceral and skeletal abnormalities and an increased risk of cancer (Cano- Gauci et al., 1999; Pilia et al., 1996; Tenorio et al., 2014; Veugelers et al., 2000). GPC3 is also a known regulator of apoptosis (Grisaru et al., 2001; Liu et al., 2012; Miao et al., 2013, 2014; Sun et al., 2011) with functions in hepatocellular carcinoma (Zheng et al., 2022). The expression of Unc5 receptors is also affected in a variety of cancers (Mehlen and Guenebeaud, 2010), and of prognostic value in neuroblastoma, where it drives tumour cell ability to migrate and/or survive (Delloye-Bourgeois et al., 2009; Wang et al., 2014) . However, most of the molecular mechanisms that underpin neuroblastoma cell dissemination remain unknown. We recently developed the first relevant *in vivo* model, by grafting human neuroblastoma cells arising from the sympathoadrenal lineage of the neural crest, into the equivalent site in chick embryos (Delloye-Bourgeois et al., 2017). Using this model, we demonstrated that neuroblastoma cells exploit Semaphorin3C/Neuropilin/Plexin signaling for metastatic dissemination, and to migrate from the neural crest domain at the edge of the neural tube to sympathoadrenal derivatives, which are the sites of primary tumors in patients (Delloye-Bourgeois et al., 2017). These tumour cells also take advantage of exogenous signals released by the developing nerve ganglia, such as olfactomedin1 (Ben Amar et al. *Nature Communications*, in press). Interestingly, GPC3 is widely expressed in embryonal tumors (Ortiz et al., 2019) and was detected in a subset of aggressive neuroblastoma samples (Dong et al., 2020).

Here we present a comprehensive structural and biophysical analysis of GPC3/Unc5 complexes. Crystal structures reveal a striking GPC3:Unc5D (4:4) octameric arrangement. An unexpected, structured glycan-glycan interaction links the C-mannosylated tryptophans on the Unc5D TSP1 domain to an N-linked glycan on GPC3, the first such interaction to be described in a membrane receptor complex. Protein-protein interfaces are formed along the concave face of the GPC3 core and engage all three N-terminal domains of Unc5D. We use mutagenesis and molecular dynamics simulations to characterise these interfaces, and present mutants that no longer interact. Anti- GPC3 nanobodies either disrupt or enhance Unc5-binding. *In vitro*, we show that a repulsive cellular response is elicited by Unc5-GPC3 signalling in HeLa, N2A, SY5Y cells and cortical neurons, suggesting that the Unc5/GPC3 signalling mechanism is conserved across different cell types. In the developing mouse cortex, we find that GPC3 is expressed on apical progenitors, and acts as a ligand for Unc5D expressed on migrating neurons. The interaction impacts the speed at which neurons migrate towards the upper cortical plate layers. In our second *in vivo* example, we show that autocrine and paracrine Unc5/GPC3 signalling is essential for the collective migration of neural-crest derived neuroblastoma cells to sympathoadrenal derivatives in which they form primary tumours.

## Results

### Structures of mouse and human GPC3

We produced human GPC3 residues 32-483 (hGPC3^core^) and murine GPC3 residues 31-482 (mGPC3^core^) in GnTI-deficient HEK293 cells and crystallised the purified proteins using the vapour diffusion method. Crystals diffracted up to 2.6 and 2.9 Å resolution, respectively. The structures were solved by molecular replacement using a model based on hGPC1 (Awad et al., 2015). Human and murine GPC3 sequences are 94% identical and we find that the two structures are similar (Cα root-mean-square deviation, RMSD_Cα_ = 0.54 Å for 353 aligned atoms), Fig. 1B and S1A. Compared to the previously solved structures of fly DLP and human GPC1, which belong to the other subclass of GPC proteins, GPC3 has a more curved shape (Fig. 1C). Superposition of hGPC1 (Awad et al., 2015) and hGPC3^core^ results in an RMSD_Cα_ = 8.87 Å (for 358 aligned atoms). We modelled glycans on two predicted N-glycosylation sites (N124 and N241 in hGPC3, Fig. 1B, N123 and N240 in mGPC3) into evident electron density (Fig. S1B). Crystallographic details are summarised in Table S1.

### GPC3 is a high-affinity ligand for Unc5 receptors and forms an octameric hetero-complex

We used an unbiased enzyme-linked immunosorbent assay (ELISA) (Ozgul et al., 2019; Ranaivoson et al., 2019) to discover that Unc5 receptors are potential GPC3-binders. We confirmed the interaction using surface plasmon resonance (SPR) binding experiments with available purified ectodomains (Fig. S1C). To produce high quality complex crystals, we over- expressed proteins in HEK293 GnTI-deficient cells and mixed hGPC3^core^, mGPC3^core^ or murine GPC3 residues 31-488 (mGPC3^488^) with *rattus norvegicus* Unc5D residues 32-307 (rUnc5D^IgIgTSP^). The complexes crystallised in two different space groups (Table S1) with different crystal packing. Strikingly, all three datasets reveal an octameric assembly (Fig. 1D-I, Fig. S1D). The centre of the octamer is formed by four Unc5D molecules that are aligned in a ‘head-to-tail’ antiparallel bundle. Two GPC3 molecules wrap around each end of the Unc5 tetramer. Each GPC3 chain interacts with three Unc5 chains, forming interfaces with the Ig1/Ig2 domains of two Unc5 molecules, and with a TSP1 domain of a third Unc5 molecule (Fig. 1D-I).

We performed native mass spectrometry to assess the stoichiometry of the complex outside the crystal lattice. Wild type GPC3 protein did not give clean signal using this method. We speculated that this may be due to a mixture of cleaved and uncleaved protein in our samples, which is only partially processed at the conserved furin-like convertase site in the C-lobe. To produce a more homogenous sample for mass spectrometry, we introduced two single point mutations in the furin cleavage site (R355A and R358A). Mixed with rUnc5D^IgIgTSP^, we reveal masses corresponding to the octamer and its subfragments: 1:1 Unc5D-Gpc3 dimers (93 kDa), 2:2 tetramers (185 kDa) and the full 4:4 octamer (370 kDa) (Fig. 2A). Each peak was subjected to tandem mass spectrometry (MS/MS) to validate the peak components (Fig. S1E-G). These results support our structural conclusions.

**Figure 2:**
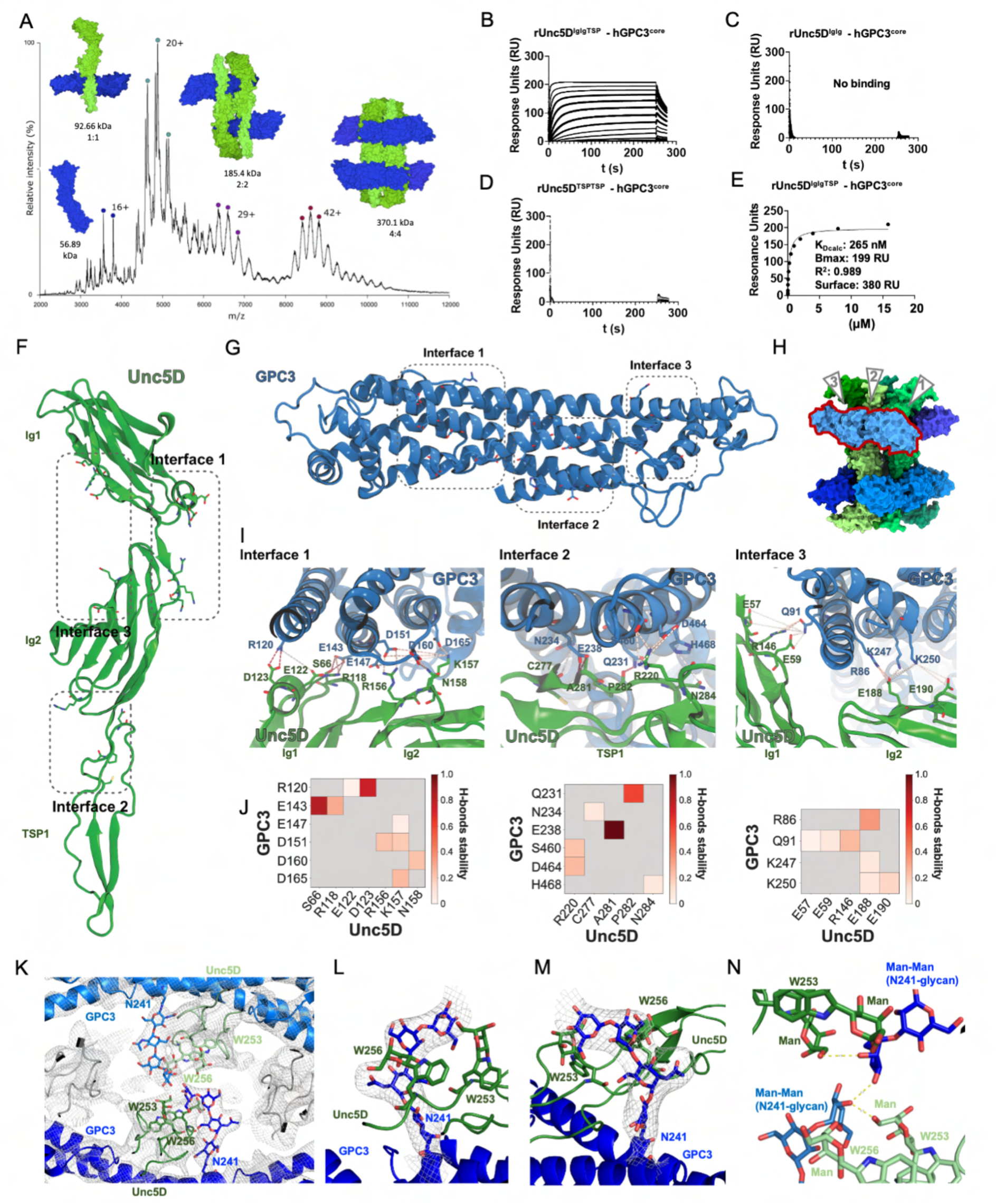
Biophysical analysis and characterization of binding surfaces in the hGPC3-rUnc5 complex. **(A)** Native MS spectrum of rUnc5^IgIgTSP^ and hGPC3^core (R355A/R358A)^. Individual peaks were isolated for MS/MS analysis to identify subcomplexes (Fig. S1E-G). **(B-E)** SPR data shows binding that rUnc5^IgIgTSP^, but not the shorter constructs rUnc5D^IgIg^ and rUnc5D^TSPTSP^, binds hGPC3^core^ with nanomolar affinity. The apparent K_D_ (K_Dcalc_) was calculated using a 1:1 binding model and is indicative only. Bmax, R^2^ and the amount of ligand immobilised on the flowcell surface are indicated. **(F)** Binding interfaces 1-3 on rUnc5D^IgIgTSP^. **(G)** Binding interfaces 1-3 on hGPC3^core^. **(H)** Binding interfaces 1-3 indicated on the octameric complex. The glypican molecule for which these are indicated is outlined in red. **(I)** Zoomed views of interacting residues in interfaces 1-3 (hGPC3^core^-rUnc5D^IgIgTSP^ complex). Hydrogen bonds are shown as dotted yellow lines. **(J)** Summaries of the hydrogen bond analyses during restrained MD simulation. Atoms that contribute to stable hydrogen bonds between the two proteins are shown, and colored blocks indicate the stability of the bond during simulation (averages for the four copies of the complex). Non-averaged results are shown in Fig. S2A-C. **(K)** View of the glycan emanating from two copies of hGPC3 N241 towards the centre of the complex. C-mannosylated tryptophans of nearby rUnc5D TSP1 domains are indicated (W253, W256). The calculated 2FoFc map of the hGPC3-rUnc5D complex data is shown as a grey mesh (sigma=1). **(L, M)** As panel K, but showing zoomed views of the N241-glycan for one of the hGPC3 copies within the complex. The map is carved around the N-linked glycan. **(N)** Distances below 3.5 Å between atoms within glycans from different chains are indicated as yellow dotted lines.

### The GPC3-Unc5 super-complex requires multiple binding surfaces

The octameric arrangement of the complex involves three main interfaces, in which the highly sequence-conserved concave surface of GPC3^core^ contacts all three N-terminal domains of Unc5: Ig1, Ig2 and TSP1 (Fig. S1H). SPR experiments show that the Ig domains alone, or the TSP domains alone, are not sufficient for detectable binding to GPC3 (Fig. 2B-E). This arrangement contrasts with that seen in Unc5-FLRT complexes, where a single interface forms between the FLRT leucine-rich repeat (LRR) domain and Unc5 Ig1 (Jackson et al., 2016; Seiradake et al., 2014). In agreement with our conclusions, the human isoform 1 of Unc5A, which lacks TSP1 and therefore incudes only Ig1, Ig2 and TSP2 (hUnc5A^iso1^), is unable to bind GPC3 (Fig. S1I). To better characterise the protein-protein binding interfaces in our model, we performed 500 nanoseconds of molecular dynamics simulations of the hGPC3^core^-rUnc5D^IgIgTSP^ complex. Stable interactions of the three main interfaces (Fig. 2F-H), averaged for the four copies within the octameric complex, are shown in Fig. 2I, J. Equivalent data for each copy is shown in Fig. S2A-C.

Interface 1 is located at the C lobe of GPC3 and interfaces with the Ig1 and Ig2 domains of Unc5D, thereby burying a surface of ∼690 Å^2^ (mGPC3 complex) or ∼640 Å^2^ (hGPC3 complex). The interacting surfaces display a high level of charge complementarity, which are contributed by hGPC3 (R120, E143, E147, D151, D160, D165) and rUnc5D (R118, E122, D123, R156, K157) (Fig. 2I, J and S2A). The largest interface, interface 2, is formed between the Unc5D TSP1 and Ig2 domains and the M lobe of GPC3, burying ∼1440 Å^2^ (mGPC3 complex) or ∼1330 Å^2^ (hGPC3 complex) of protein surface. Hydrophobic residues line both sides of the interacting surfaces: rUnc5D I170, A281, P282, L283, F288, and hGPC3 L157, L235. These are complemented by extensive hydrogen bonding and charged interactions, such as hGPC3 E238 which interacts with the backbone of rUnc5D A281 (Fig. 2I, J and S2B). Interface 3 involves the N lobe of GPC3 and contains many charged and hydrogen bonding interactions (Fig. 2I, J and S2C). The interacting surfaces are contributed by two distinct patches on Unc5D that are located on the Ig1 and Ig2 domains. The total buried surface per interface is ∼390 Å^2^ (mGPC3 complex) or ∼360 Å^2^ (hGPC3 complex). Within the compact octameric arrangement, the antiparallel lying Unc5D chains form extensive interactions between themselves, especially at the Ig2 domains: the buried surface amounts to ∼900 Å^2^ and ∼840 Å^2^ for each of the antiparallel packing interactions within the Unc5 bundle.

### The Unc5-GPC3 complex is stabilised by an essential inter-chain glycan interaction

Recent work showed that the TSP domains of Unc5 receptors are C-mannosylated at tryptophan residues W_1_ and W_2_ of the consensus sequence W_1_xxW_2_xxW_3_ (Shcherbakova et al., 2019). We revisited our 2.4 Å-resolution published crystallographic data for hUnc5A^iso^ (Seiradake et al., 2014) and noticed evidence for the presence of C-mannosylation (Trp245 and Trp248), Fig S2D, E. We purified hUnc5A^iso1^, hUnc5B^IgIgTSP^ (residues 26-303), mUnc5C^IgIgTSP^ (residues 40-317), and rUnc5D^IgIgTSP^ from HEK293T cells and performed mass spectrometry to verify that C-mannosylation is indeed present on these proteins. The results show that the first two tryptophan residues (W_1_ and W_2_) were C-mannosylated in all Unc5 homologues tested (Fig. S2F-I). In agreement with these results, we observed electron density for these glycans in the crystallographic maps calculated for the rUnc5D complexes described above.

N-linked glycan chains are flexible and usually not defined in crystal structures unless they are held in place by specific interactions. The electron density maps calculated for the GPC3-Unc5D complexes reveal extra density extending from hGPC3 N241, one of the predicted N-linked glycosylation sites, towards the centre of the complex. They pack closely against the C-mannosylated tryptophans 253 and 256 of the rUnc5D TSP1 domain (Fig. 2K-N). In the unliganded structures of human and mouse GPC3 this glycan chain is not visible. We sought to remove this glycan *in vitro*, to test its function. Attempts to remove the hGPC3 N241-linked glycan using endoglycosidases (EndoF1 or PNGase F), with the aim to assess its role in Unc5 binding, were not successful, as the glycan remained uncleaved. We therefore mutated the glycosylation site (N241Q) to produce a hGPC3 protein that lacks the glycan at this position. The resulting mutant protein was readily expressed and secreted by HEK293 cells. We used a cell-based binding assay to test whether this mutant protein still binds to mUnc5B expressed on the surface of HEK293T cells and found no binding (Fig. 3A,B). SPR experiments confirm these results (Fig. 3C) and show that the N241Q mutant has lost affinity also for other Unc5 receptors (Figs. 3D and S3A, B). To be consistent with previously used nomenclature we will be referring to this mutant as GPC3^UG^, with UG standing for ‘non-Unc5-binding GPC3’, throughout the rest of this manuscript. To produce non-GPC3 mutants of Unc5, we used an established approach where an artificial N-linked glycosylation site is engineered to disrupt protein-interaction (Jackson et al., 2015, 2016, 2018; Seiradake et al., 2013, 2014; del Toro et al., 2020) (Fig 3B). These Unc5 mutants contain a mutation in binding interface 2 (A277N+L279T in hUnc5B) and they still bind the canonical ligand FLRT2, but not GPC3. In analogy to previous nomenclature, we call the resulting non-GPC3 binding mutants: Unc5^GU^.

**Fig. 3.**
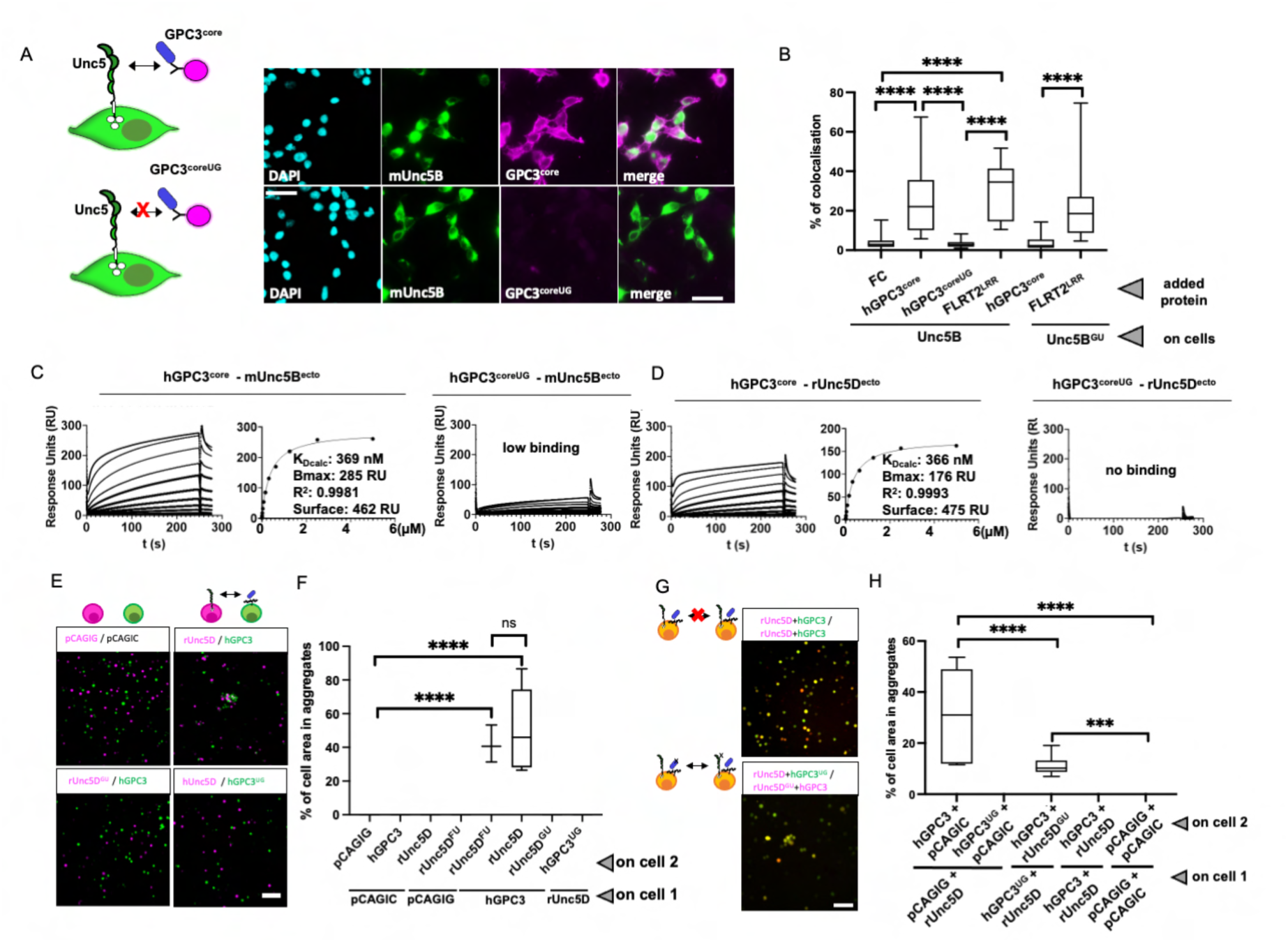
The non-binding mutants Unc5^GU^ and GPC3^UG^ display impaired GPC3-Unc5 binding, Unc5-GPC3 interaction ‘*in trans’* is inhibited by ‘*in cis’* interactions. (A) A cell-based binding assay shows binding between mUnc5B (expressed on cells) and purified GPC3^core^ protein, visualised by immunostaining of the poly-histidine tag. The N241Q (GPC3^coreUG^) mutant protein does not bind. Representative images are shown. (B) Quantification of experiments shown in panel A, and of additional conditions tested. The mUnc5B A277N+L279T (mUnc5B^GU^) mutant binds the canonical ligand FLRT2 (LRR domain, FLRT2^LRR^), but not hGPC3^core^. (C) SPR data confirms that hGPC3^coreUG^ has strongly reduced affinity to Unc5B^ecto^ compared to wild type protein. The apparent K_D_ (K_Dcalc_) was calculated using a 1:1 binding model and is indicative only. Bmax, R^2^ and the amount of ligand immobilised on the flowcell surface are indicated. (D) As panel C, using rUnc5D^ecto^. (E) A cell-cell aggregation assay shows that GPC3 and Unc5D mediate cell adhesion in *trans.* Representative images are shown. (F) Quantification of the experiments described in panel E, using single transfected cells. Empty vector controls: pCAGIC (red) and pCAGIG (green). Wild type and mutant receptors were used: non-FLRT-binding mutant: rUnc5D^FU^, non-GPC3-binding mutant: rUnc5D^GU^, non-Unc5-binding mutant: hGPC3^UG^. (G) Co-expression of Unc5D and GPC3 *in cis* interferes with *in trans* interaction and cell-cell aggregation. Representative images are shown. (H) Quantification of the experiments described in panel G, using double transfected cells. *** p<0.001; **** p<0.0001, one-way ANOVA with Tukey’s post hoc tests. Scale bars represent 100 µm.

### GPC3-Unc5 binding promotes cell-cell *‘in trans’* interaction

The intriguing geometry of the octameric complex begs the question whether these proteins interact on the surface of the same cells ‘*in cis’* or across cells ‘*in trans’*. We used an established cell aggregation assay (Berns et al., 2018; Pederick et al., 2018, 2021; del Toro et al., 2020). The protein constructs used in this assay are ‘full length’ versions and anchored at the cell surface. Aggregation in these assays is demonstrative of an *in-trans* interaction. We found that Unc5D- expressing K562 cells bind and aggregate with GPC3-expressing cells *in vitro* (Fig. 3E, F). The cells did not aggregate when the wild type proteins were replaced with either of the ‘non-binding‘ mutants, rUnc5D^GU^ and hGPC3^UG^. Conversely, the non-FLRT binding rUnc5D (rUnc5D^FU^, (Seiradake et al., 2014)) still causes aggregation with GPC3-expressing cells, confirming that the affected binding sites are distinct (Fig. 3E, F), as supported also by the structural data. We also co-expressed Unc5D and GPC3 on the same population of cells to test whether *in cis* binding interferes with the ‘*trans’* interaction described above, as seen for other receptors such as Eph/ephrins (Carvalho et al., 2006). Indeed, we found that cells co-expressing rUnc5D and hGPC3 did not aggregate (Fig 3G, H), indicating that *in cis*-interaction silences ‘*trans’* binding. In agreement with this finding, the co-expression of WT rUnc5D + hGPC3^UG^ on one cell population, and WT hGPC3 + rUnc5D^GU^ on the other population led to aggregation, showing that *in cis* interaction, rather than just co-expression, is required for silencing. *In cis* silencing can occur due to sequestering of binding surfaces on the cell surface, or due to inhibition of cell surface presentation of complexed proteins. We quantified the expression of the receptors using western blot analysis to reveal total protein, and by surface immunostaining to reveal cell surface presentation. The results demonstrate that co-expression does not prevent surface presentation of the receptors (Fig. S3C-G). Taken together, the data suggest a mechanism by which the interaction can occur both *in cis* and *in trans*. However, when proteins are co-expressed in the same cell, then *in cis* silencing of *trans* interaction occurs by occupying the available interaction sites.

### Characterisation of functional anti-GPC3 nanobodies: Nano^glue^ and Nano^break^

To generate additional tools for the functional characterisation of the interaction, we characterised two llama-derived heavy-chain chain antibody-derived nanobodies, which bind to murine and human GPC3^core^ (Fig. 4A-C). Pull-down data suggests that one nanobody enhances complex formation between GPC3 and Unc5 (Nano^glue^), whilst another inhibits it (Nano^break^) (Fig. 4D). We confirmed these results using SPR experiments, where the nanobodies enhance or compete with Unc5-GPC3 binding (Fig. 4E,F, Fig. S4A-E). The pulldown and SPR results also show that Nano^break^ has an overall weaker affinity compared to Nano^glue^. Consistent with the protein-binding studies, our cell aggregation assays show that the addition of nanobody Nano^break^, and not Nano^glue^, inhibited Unc5D-GPC3 mediated cell-cell adhesion (Fig. 4G, H). Of note is that we did not observe enhanced aggregation with Nano^glue^ in this assay, which may be due to the strong aggregation phenotype observed also in absence of the nanobody.

**Figure 4.**
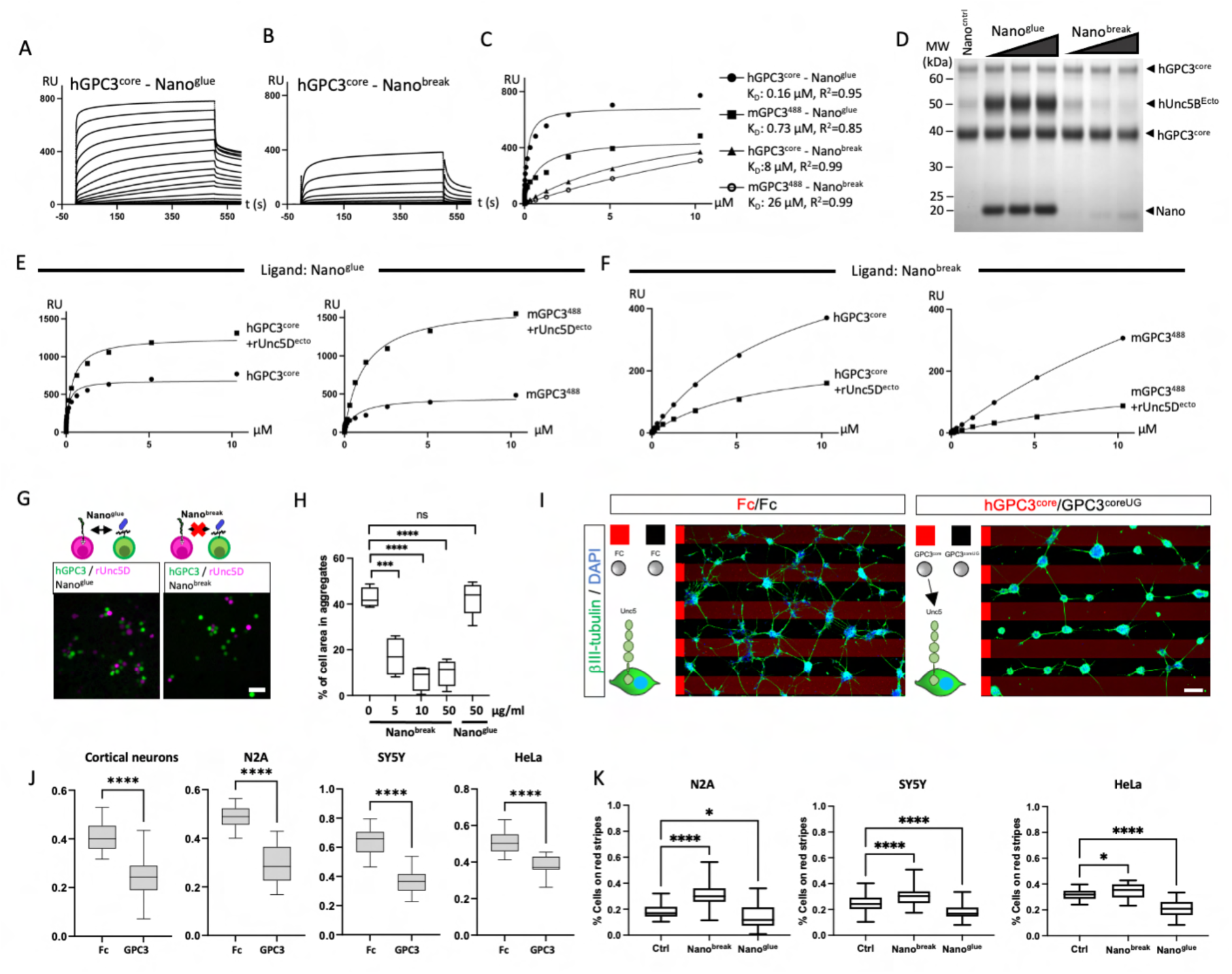
Nanobodies enhance or disrupt GPC3-Unc5 interaction. **(A)** SPR binding curves are plotted for different concentrations of hGPC3^core^ injected over immobilized Nano^glue^. **(B)** As in panel A, with Nano^break^. **(C)** The equilibrium values from experiments shown in panels A and B, and equivalent values from using mGPC3^488^ (Fig. S4D, E), were plotted. K_D_ values are calculated assuming 1:1 binding. **(D)** Pull-downs suggest that Nano^glue^ enhances Unc5B- binding to immobilised hGPC3^core^, whilst Nano^break^ weakens the interaction. **(E,F)** Equilibrium SPR data (see curves in Fig. S4D, E) confirms that Nano^glue^ enhances, and Nano^break^ weakens, GPC3 - Unc5D binding. **(G)** We used a cell aggregation assay to show that the addition of Nano^break^ inhibits GPC3 - Unc5D cell adhesion in *trans.* Nano^glue^ does not interfere with Unc5-GPC3-mediated cell aggregation in these experiments. Representative images are shown. **(H)** Quantification of the cell aggregation experiments described in panel G. **(I)** E15.5 dissociated cortical neurons were grown on alternate stripes (red and black) containing Fc, GPC3^core^ or GPC3^coreUG^. Neurons were stained with beta-III-tubulin (green) and DAPI (blue). **(J)** Quantification of the percentage of beta-III-tubulin (green) pixels on red stripes: for different experiments Fc (=Fc/Fc control), GPC3 (=hGPC3^core^/hGPC3^coreUG^). We performed equivalent stripe assays also for HeLa, N2A and SY5Y cells, using DAPI for quantification. Student T-tests were performed. **** p<0.0001. **(K)** We performed stripe assays as shown in panel I, but in the presence of streptavidin (Ctrl) or streptavidin-nanobody complexes (Nano^glue^ or Nano^break^). In agreement with enhancing or disrupting the Unc5-GPC3 interaction, we found that Nano^glue^ and Nano^break^, enhanced or reduced the repulsive response from wild type GPC3^core^, respectively, compared to Ctrl. We performed one-way ANOVA with Tukey’s post hoc tests. NS, not significant; * p<0.05; *** p<0.001; **** p<0.0001. Scale bars represent 100 μm (G) and 90μm (I).

### GPC3-Unc5 interaction produces contact-repulsion *in vitro*

Unc5 receptors and GPC3 play key roles in many tissues. In particular, Unc5 receptors are known for their repulsive signalling in neuronal cell guidance (Round and Stein, 2007; Yamagishi et al., 2011). To assess if GPC3-Unc5 interaction is mediating contact-repulsion, we used stripe assays and tested the response of selected cell types known to express Unc5 receptors (Delloye-Bourgeois et al., 2009; Mehlen and Guenebeaud, 2010; Seiradake et al., 2014; Wang et al., 2014). Cells were plated on alternating stripes of purified GPC3^core^ and the mutant GPC3^coreUG^. In these assays, cortical neurons preferentially migrated on the mutant protein stripes, demonstrating that GPC3^core^ elicits a repulsive effect via an Unc5-dependent mechanism (Fig. 4I, J). Interestingly, when given the choice between GPC3^core^ (WT or mutant) and neutral Fc protein, the neurons were strongly repelled by both wild type and the mutant protein, suggesting that unknown additional GPC3-receptors, who do not depend on the Unc5- GPC3 interaction, also cause repulsion from GPC3^core^ (Fig. S4F-G). Unc5 receptors and GPC3 play key roles in many tissues, and therefore the mechanisms revealed here could be of importance in other cell types too. We then tested HeLa, N2A and SY5Y neuroblastoma cell lines in stripe assays using alternating stripes of WT GPC3^core^ and the mutant GPC3^coreUG^. As observed for the neurons, all of these cells exhibited a preference for the mutant protein (Fig. 4J). We also tested the effect of the nanobodies in these stripe assays. We tetramerised the nanobodies via a biotinylated linker and streptavidin to increase their affinity and potency. For N2A, SY5Y and HeLa cells, the addition of nanobodies tended to enhance (Nano^glue^) or decrease (Nano^break^) cell repulsion from wild-type GPC3-containing stripes (Fig. 4K). We also attempted to perform this assay with cortical neurons, however the addition of Nano^glue^ led to immobilisation of these cells on the stripe surface, and we were therefore unable to quantify any migratory behaviour (not shown). Conversely, we found that addition of Nano^break^ reduced the repulsive effect of wild type GPC3^core^ (Fig. S4H-I) as observed for other cell types. We conclude that the GPC3- Unc5 interaction mediates contact-repulsion in these assays, and may contribute to cellular navigation during the migration of cortical and neuroblastoma cells.

### GPC3 and Unc5D are expressed in the developing mouse brain cortex

We previously showed that Unc5D is expressed in the cortex during development where it regulates radial migration (Seiradake et al., 2014; Yamagishi et al., 2011) . Glypicans also show specific patterns of expression during central nervous system development, with five out of six known glypicans (GPC1-4 and GPC6) expressed at earlier stages of brain development (Ford-Perriss et al., 2003) and in neural stem cells (Oikari et al., 2016). Here we asked whether Unc5D- GPC3 could regulate cortical migration. *In situ* hybridization (ISH) for GPC3 showed restricted expression to the germinal layers, predominantly at the ventricular zone (VZ) where AP cells are located from embryonic days 13.5 [E13.5] to E17.5 (Fig. 5A, B and S5A, B). Unc5D showed strong expression in areas enriched in young/migrating neurons (SVZ/IZ) as reported previously (Miyoshi and Fishell, 2012; Takemoto et al., 2011).These results were confirmed by co-staining with the neuronal marker Ctip2 and the apical progenitor marker Pvim (Fig. 5C). Analysis using data from two single-cell RNA seq databases showed that Unc5D is enriched in migrating neurons, while GPC3 is expressed predominantly in apical progenitors (AP) of the mouse cortex, from E13.5 to E17.5 (Fig. 5D and (Florio et al., 2015), Fig. 5E-F, S5C-D and (di Bella et al., 2021)). Distribution analysis using categorized clusters showed that 66% of Unc5D-positive cells are migrating neurons, whereas 57% of GPC3+ cells are apical progenitors at E15.5 (Fig. 5G). Pull-down experiments using Nano^glue^ in E15.5 mouse cortex lysate showed high specificity and enrichment of GPC3 protein. Moreover, Unc5D co-immunoprecipitated with GPC3, suggesting that the two proteins interact *in vivo* (Fig. 5H-I). Consistent with the expression data, we found that GPC3 protein is present in the germinal zone, mainly where the AP cell bodies are located within the VZ, and to a lower extend in the IZ and CP, where the pattern resembles that of AP fibers and their end feet (Fig. 5J). Based on these results, we developed a working model in which migrating neurons expressing Unc5D interact with GPC3 present in AP cells (Fig. 5K).

**Figure 5.**
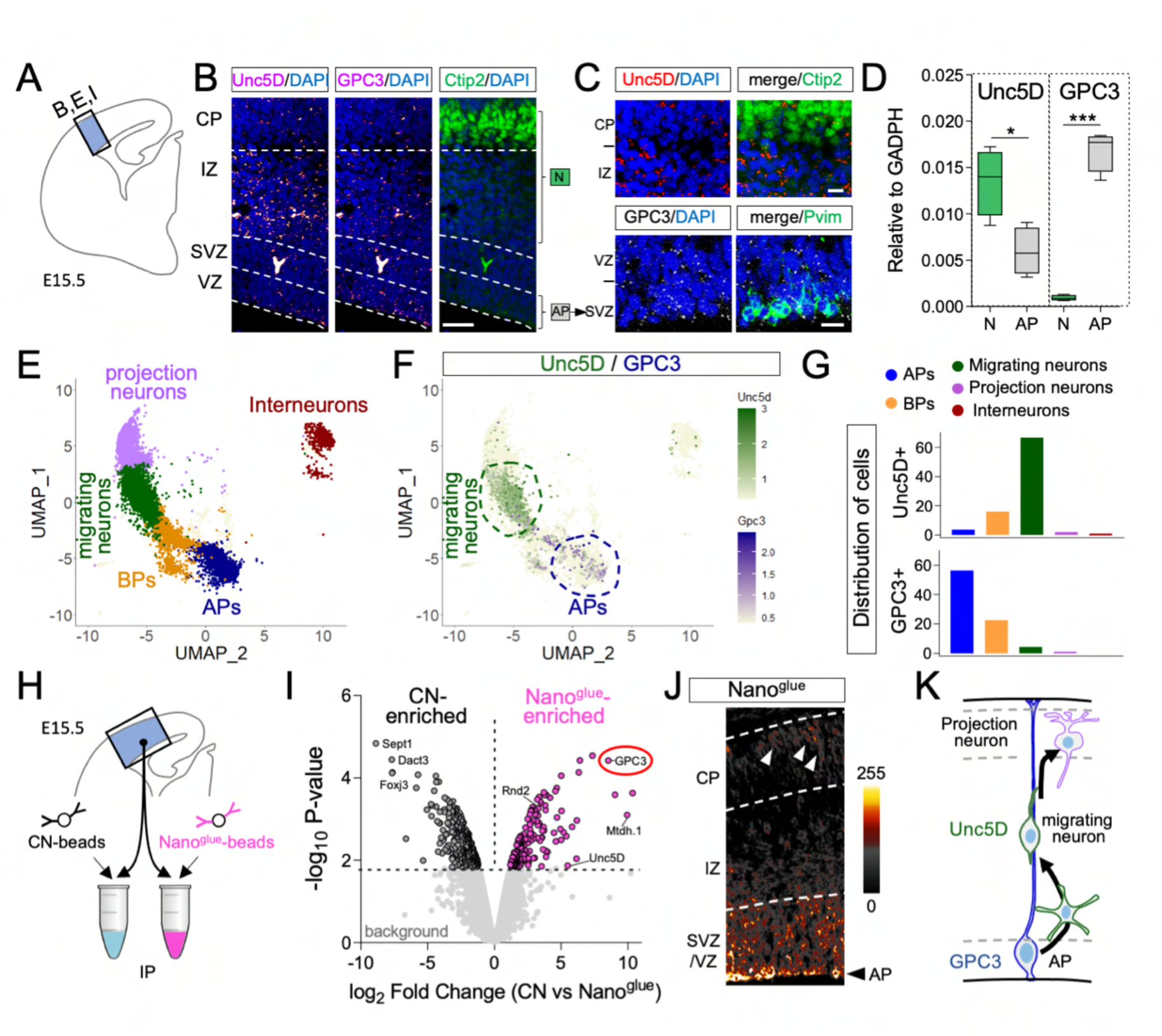
GPC3 is expressed by cortical apical progenitor cells and interacts with Unc5D. **(A)** Cortical region shown in B, J and magnified in C. **(B)** *In situ* hybridization (ISH) for Unc5D and GPC3 (magenta) shows their expression in the cortex at E15.5. Nuclear staining with DAPI is shown in blue. The layers enriched in neurons (N) and apical progenitors (AP) are indicated. **(C)** Upper panels show the ISH for Unc5D (red) combined with the neuronal marker Ctip2 (green). Lower panels show the ISH for GPC3 (white) and the apical progenitor marker Pvim (green). The location of the CP, IZ, SVZ and VZ layers are indicated. **(D)** Unc5D and GPC3 expression levels normalized to GADPH in neurons and APs, using RNA profiling data published in Florio et al., 2015 (GSE65000). Unc5D mRNA is high in neurons (N), while GPC3 mRNA is enriched in apical progenitors (AP). *p < 0.05, ***p<0.001, two-tailed Student’s t test. **(E)** UMAP visualization of single-cell data from E15.5 mouse cortex published in Di Belal et al., 2021. Five major cell clusters are colored by cell-type assignment based on published metadata (GSE153164) (basal progenitor, BP, apical progenitor AP). **(F)** Combined plot of Unc5D (green) and GPC3 (magenta) mRNA expression per cell. Most of Unc5D-expressing cells belong to the migrating neuron cluster (dashed green line), while GPC3-expressing cells are highly enriched in the apical progenitor (AP) cluster (blue dashed line). **(G)** Quantification of the distribution of Unc5D- and GPC3-positive cells across all five major clusters. **(H)** Cortical region used for pull-down with Nano^glue^. **(I)** Volcano plot showing enriched proteins in control and Nano^glue^ pull-downs revealed by mass spectrometry using label-free quantification (LFQ). Proteins enriched in the Nano^glue^ pull-down are colored in pink and those enriched in the control condition (streptavidin agarose beads) in black. Non- significant proteins are colored in gray and labelled as background. **(J)** Immunostaining for GPC3 using Nano^glue^ coupled to fluorescent streptavidin on coronal section of E15.5 mouse cortex. The image is colored based on the intensity of the staining and shows high expression in the germinal zones, mainly at the apical progenitor (AP) cell bodies. Arrowheads indicate staining that resembles the pattern of AP fibers in the CP. **(K)** Working model showing that migrating neurons expressing Unc5D (green) could interact with GPC3 present in AP fibers (blue). Scale bars represent 200 μm (B, J) and 20 μm (C).

### GPC3-Unc5 interaction is required for radial neuronal migration *in vivo*

To study the effects of Unc5D-GPC3 binding on cortical neuron migration, we used in-utero electroporation at E13.5 to reveal potential functions of GPC3 in cortical migration during development. We previously showed that the over-expression of signalling-deficient, but otherwise active, receptor fragments is an effective way of interfering with endogenous interactions (del Toro et al., 2020). Here, we overexpressed rUnc5D^IgIgTSP^ in migrating neurons to study the effect of depleting Unc5-binding sites on GPC3 (Fig. 6A), by competing with endogenous Unc5 receptors. Consistent with the role of Unc5D in radial migration, expression of a secreted version of its ectodomain produced a strong delay in neuronal migration (Fig. 6A- B). The accumulation observed in the IZ resembles the phenotype seen when full length Unc5D is overexpressed in migrating neurons (Seiradake et al., 2014; Yamagishi et al., 2011). This effect was partially rescued when using the mutant Unc5D^IgIgTSPGU^ (Fig. 6A-B, Fig. S6A), confirming that the migration delay is at least partially due to interaction with GPC3. In an alternative approach to reduce interactions, we knocked down endogenous GPC3 in E13.5 cortices using the small hairpin RNA (shRNA) target sequence in the pCAG-miR30 vector system (Matsuda and Cepko, 2007) (Fig. S6B). We used the pCAG-BLBP vector to visualize the targeted apical progenitors and their fibers and measured the distribution of wild type neurons labelled with a mCherry reporter (Fig. 6C). Analysis at E16.5 showed reduced migration of neurons along the fibers deficient for endogenous GPC3 protein, supporting a key functional role (Fig. 6C, D). We also over-expressed secreted Nano^break^ and Nano^glue^ in migrating neurons (Fig. S6C), to reveal the effects on migration of the nanobodies *in vivo*. Both Nano^break^ and Nano^glue^ overexpression caused significant delays in migration to the upper cortical plate (Fig. 6E, F). Taken together, these results show that Unc5-GPC3 interactions regulate cortical migration *in vivo*.

**Figure 6.**
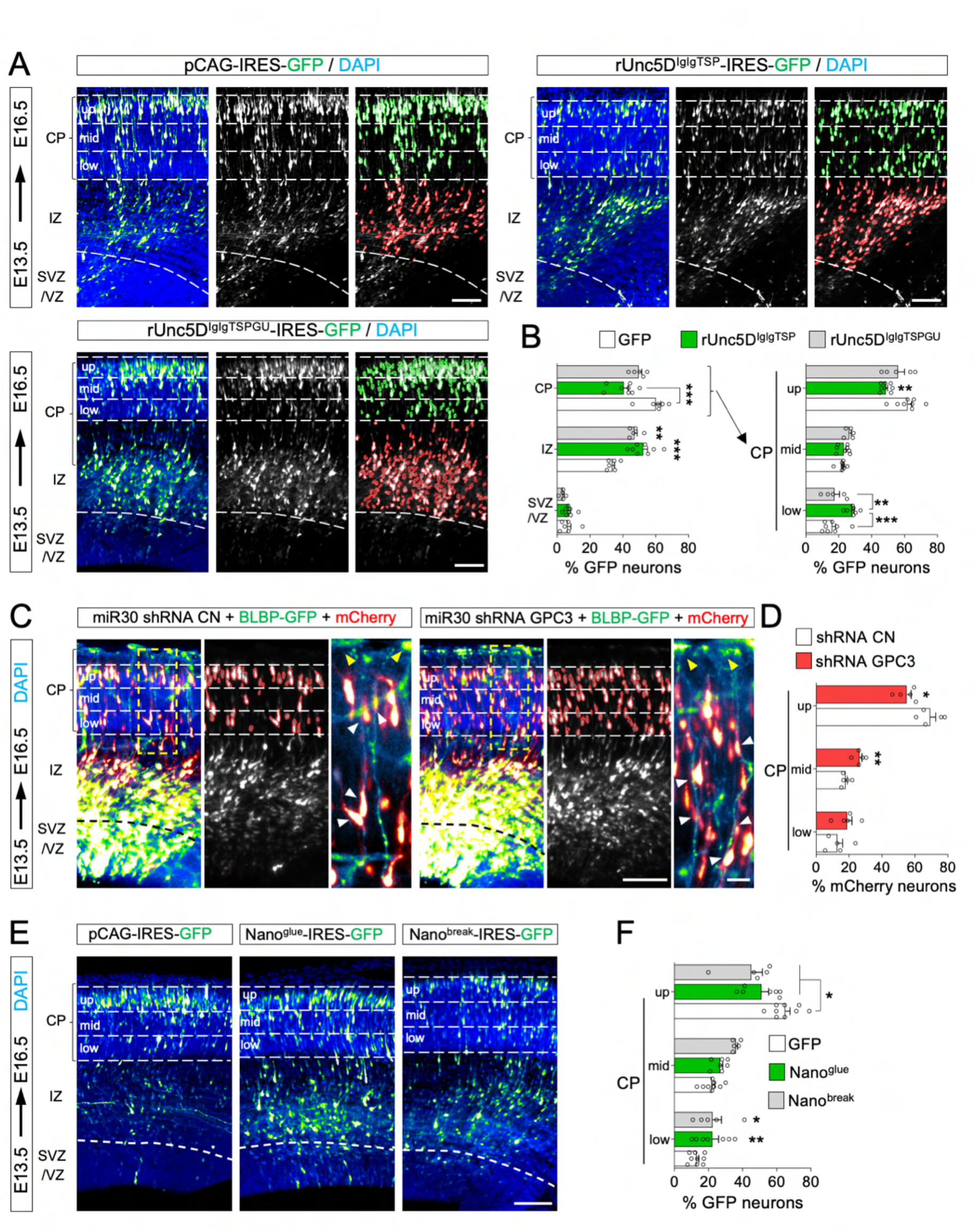
GPC3 promotes radial migration of Unc5-expressing cells. **(A)** Coronal sections of E16.5 cortex after *in utero* electroporation (IUE) of empty pCAG- IRES-GFP (pCAGIG, control), or pCAGIG encoding rUnc5D^IgIgTSP^ or rUnc5D^IgIgTSPGU^ at E13.5. The cortical plate (CP) is defined based on the DAPI staining. GFP-positive cells in the IZ and CP are automatically identified (outlined in green and red respectively) and the percentage in each layer is quantified. The CP is further subdivided into 3 bins (up, mid, and low) to analyse the distribution of GFP-positive cells in each bin. **(B)** Quantification of data shown in (A). n = 8 GFP, n = 8 rUnc5D^ecto^, and n = 5 rUnc5D^IgIgTSP^ electroporated brains. **p < 0.01, ***p < 0.001, one-way ANOVA test with Tukey’s post hoc analysis. **(C)** Coronal sections of a E16.5 cortex electroporated with pCAG-mCherry, pCAG-BLBP-GFP and a pCAG-miR30 vector coding for shRNA control (CN) or shRNA targeting murine GPC3. The CP was subdivided into 3 bins (up, mid, and low), and the number of mCherry-positive neurons in contact with a GFP-positive radial fiber in each bin was quantified (white arrows, inset on the right). **(D)** Quantification of the data shown in (C). n = 5 CN, n = 5 shRNA GPC3, electroporated brains. *p < 0.05, **p < 0.01, two-tailed Student’s t test. **(E)** Coronal sections of E16.5 cortex after IUE using empty pCAGIG (control), or pCAGIG encoding Nano^glue^ or Nano^break^ at E13.5. GFP- positive neurons localized within the CP were quantified for each bin. **(F)** Quantification of data shown in panel E. n = 9 GFP, n = 7 Nano^glue^, and n = 5 Nano^break^ electroporated brains. *p < 0.05, **p < 0.01, one-way ANOVA test with Tukey’s post hoc analysis. Scale bars represent 100 μm (A,C,E) and 20 μm (inset in C).

### GPC3-Unc5 interaction is required for neuroblastoma cell migration *in vivo*

The phenotypes observed in cortical migration, together with the widely documented roles of Unc5 and GPC3 in cancer, led us to investigate whether Unc5/GPC3 interaction plays a role in neuroblastoma cell migration. We analyzed the expression of GPC3 and Unc5 receptors in published single cell RNASeq data from 16 different neuroblastoma patient samples (Dong et al., 2020). Unsupervised clustering of patients’ cell data led to a segregation of tumor cells from those of the stroma: endothelial cells, fibroblasts, myeloid cells and immune cells (Fig. 7A). Interestingly, the fraction of tumor cells expressing at least one Unc5 receptors was higher in the tumor cell cluster as compared to fibroblasts, immune or myeloid cells clusters (Fig. 7B, C). Endothelial cells also highly express Unc5 receptors, especially Unc5B, as shown by others (Larrivée et al., 2007; Lu et al., 2004). Conversely, while GPC3 was also detected in a fraction of tumor cells, its expression was more frequent (29%) in fibroblastic cells of the tumor microenvironment (Fig. 7B, C). Next, we assessed GPC3 and Unc5 expression in a set of human neuroblastoma cell lines (Fig. 7D). As found for the patient tumor cells, neuroblastoma cell lines also expressed Unc5 receptors. Low GPC3 expression was detected for two out of four cell lines, hence mimicking its low and heterogeneous expression in patient samples. We selected the SY5Y cell line to further study potential roles of GPC3-Unc5 interactions in neuroblastoma cell migration. Human-specific GPC3 siRNA reduced the expression of GPC3 in these cells by 71% (± 11%, 48 hours after transfection) (Fig. S7A). In a transwell assay, the GPC3 siRNA-transfected cells showed a significant decrease in their migratory abilities compared to mock-transfected cells (Fig. S7B). These results suggest a role of GPC3-Unc5 signalling in these cells. SY5Y cells also readily overexpress transfected constructs, such as our secreted nanobodies (Fig. S7C). We used our previously established *in vivo* model of neuroblastoma (Delloye-Bourgeois et al., 2017) to perform xenografts (Fig. S7D, 7E-H). In this avian host model, neuroblastoma cells are engrafted within the pre-migratory trunk neural crest, and then migrate following a neural-crest like stereotypical ventral migratory path to the developing sympathetic ganglia and adrenal medulla (Delloye-Bourgeois et al., 2017). There, they express characteristic tumor features, forming tumor masses before undergoing secondary metastatic-like dissemination. Compared to embryos engrafted with scramble siRNA-transfected SY5Y, GPC3 siRNA-transfected cells formed tumor masses almost exclusively outside the sympatho-adrenal territories (Fig. 7E, G). In addition to being mistargeted, a high proportion of cells were dispersed and no longer integrated in the collective migration flow. These isolated cells were either delayed within the stereotyped ventral migratory route, or mislocated outside of the ventral neural crest stream. The results suggest that interfering with the neuroblastomal source of GPC3 disrupts cell migratory capacities and targeting to the primary tumor site. Next, we performed graft experiments with SY5Y cells transfected either with the vector encoding Nano^break^ or Nano^glue^ (Fig. 7F, H). Using Nano^break^, we found that manipulation of GPC3/Unc5 interactions significantly modified the migratory pattern of neuroblastoma cells: only a few isolated cells reached the sympatho-adrenal target derivatives, with most tumor masses formed outside. This phenotype is comparable to that found in the siRNA knock-down experiments described above. Interestingly, not only the nanobody-transfected SY5Y cells exhibited abnormal migratory and targeting patterns, but also the untransfected cells present in these grafting experiments. This suggests that Nano^break^ has both autocrine and paracrine effects on the collectivity of migrating neuroblastoma cells. Conversely, we found that grafted cells expressing Nano^glue^ had “enhanced” migratory properties, resulting in a preferential localization of tumor masses in the most distal trunk neural crest territories, below the dorsal aorta, in the developing adrenal gland and in enteric ganglia. This was also reflected by the smaller number of cells that failed to reach the primary tumor site, and that these tumors were highly condensed. These results agree with the SY5Y *in vitro* stripe results presented in Fig. 4K, where Nano^glue^ enhances the Unc5-GPC3 dependent cell response, whilst Nano^break^ reduces it. Taken together, the results show that modifying the strength of GPC3-Unc5 interaction determines cancer cell migration properties and tumour targeting, in the model presented.

**Figure 7:**
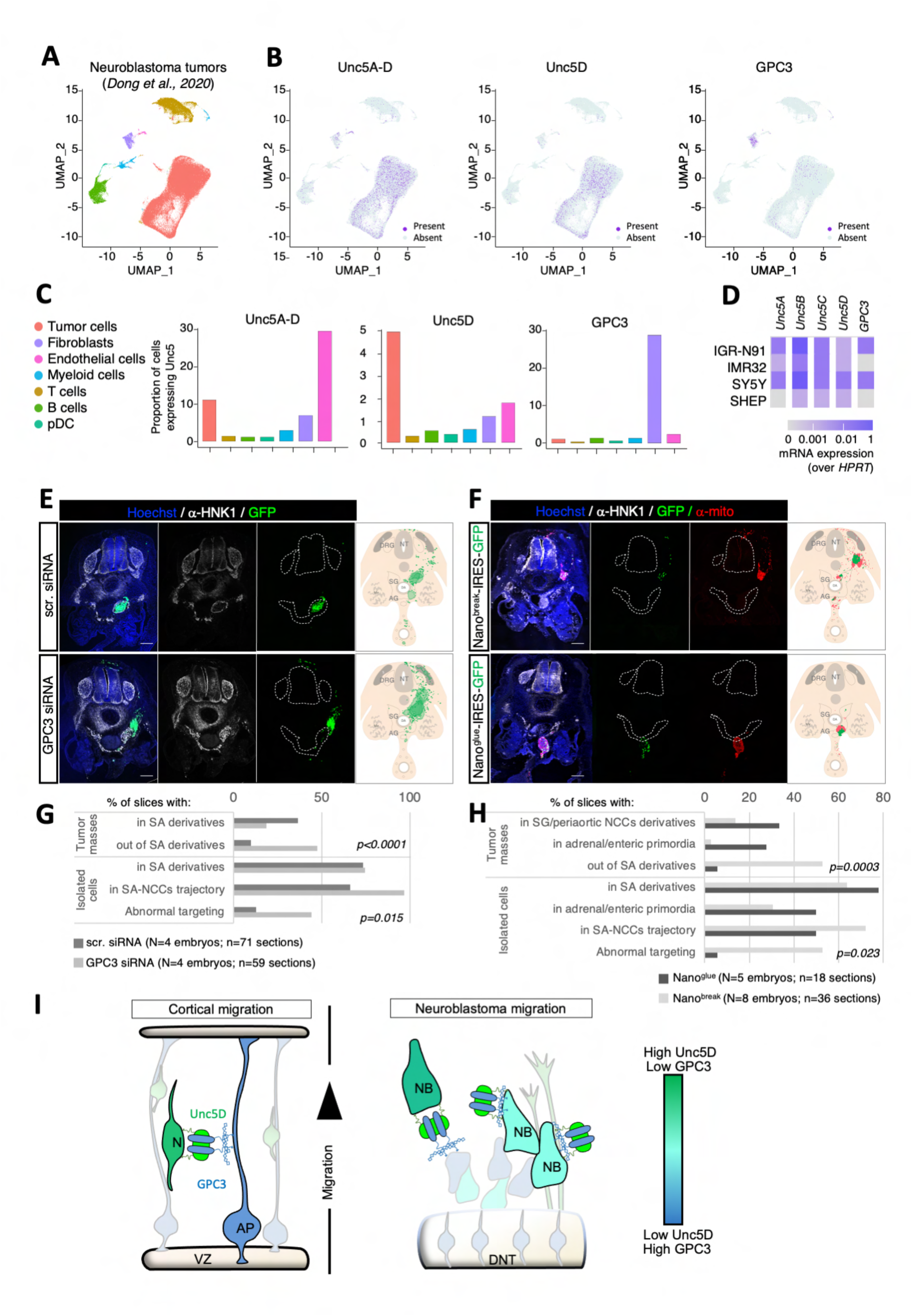
GPC3-Unc5 signalling determines neuroblastoma cell migration properties. **(A)** UMAP visualization of single-cell data from neuroblastoma tumors (Dong et al., 2020). Quantification for Unc5A-D, Unc5D alone, and GPC3 is shown in panel B. (**B**) Feature plots for Unc5A-D, Unc5D and GPC3. **(C)** Quantification of the data shown in panel B, for each cell type. **(D)** Heatmap of Unc5A-D and GPC3 mRNA expression in 4 human neuroblastoma cell lines, measured by Q-RT-PCR. (**E,F**) SY5Y cells were grafted within the migratory trunk neural crest of E2 chicken embryos and tumor positions identified 2 days later. Previous to the graft, SY5Y cells were transfected with either scr control or GPC3 siRNAs (panel E) or pCAGIG vectors encoding Nano^glue^ or Nano^break^ (panel F). Neural crest-derived structures were labeled with an anti-HNK1 antibody. Nuclei were stained with Hoechst. In E, SY5Y:GFP (green cells) are shown. In F, transfected SY5Y cells were also labeled with an anti-human mitochondrial antibody to reveal both transfected (green and red) and non-transfected (only red) SY5Y cells. Scale bar: 200 µm**. (G, H)** Quantification of SY5Y cell and tumor positions two days after grafting. We used χ^2^ tests to compare scr. (control) versus GPC3 siRNA conditions (panel E), and Nano^glue^ versus Nano^break^ conditions (panel F). NT: Neural Tube; S: Somite; DRG: Dorsal Root Ganglia; SG: Sympathetic Ganglia; AG: Adrenal Gland; DA: Dorsal Aorta; Me: Mesonephros. **(I)** Schematic summarizing the Unc5/GPC3 expression levels and putative interactions in the cortical and neuroblastoma models presented in this manuscript. Expression of Unc5D and GPC3 is color-coded from green (high Unc5D/ low GPC3) to blue (low Unc5D, high GPC3). Cells colored in cyan indicate co-expression of both receptors. N: neuron, AP: apical progenitor, VZ: ventricular zone, NB: neuroblastoma cell, DNT: dorsal neural tube. Relevant structural models are discussed in Fig. S7H, I.

## Discussion

Individual receptor-ligand interactions are embedded within complicated cell surface interactomes where most receptors bind multiple ligands. A variety of complexes are therefore formed, depending on which binding partners are available. They thereby drive many different context-dependent signalling pathways and cellular responses. This structural/functional complexity has hampered progress with understanding where specific signalling interactions act *in vivo*. Here we have employed an integrated structure-function approach that uses engineered mutant proteins and specific nanobodies to focus on a novel interaction between Unc5 receptors and GPC3. The structural data shows that these proteins form an unexpectedly large multimeric complex, with four copies of each molecule arranged in a pseudo-symmetric arrangement. Interestingly, multimeric extracellular complexes that go beyond simple (1:1) receptor:ligand interactions are emerging for a range of important morphogen and guidance receptors, for example, FLRT/Lphn/Teneurin (del Toro et al., 2020), netrin/RGM/neogenin (Robinson et al., 2021), Eph/ephrin (Seiradake et al., 2010, 2013), semaphorin/plexin/neuropilin (Janssen et al., 2012). This study and previous data show that Unc5 engages in multiple different complexes (with netrin, GPC3, with FLRT, with FLRT/Lphn), pointing to a picture where a balance between different signalling configurations is dictated by the molecular composition of the environment. Interestingly, we found that, under the harsh conditions of a native mass spectrometry experiment, the GPC3/Unc5 octamer partially disintegrates into smaller subcomponents that include 2:2 and 1:1 complexes, suggesting that these presumably weaker assemblies could also form, perhaps at initial stages of complex formation (Fig. S1E-G and Fig. 2A). Future work could address whether a full octamer is required for *cis/trans* formation of the complex, for example, using advanced electron tomography. The structures also reveal certain unexpected and striking similarities. For example, the tight anti-parallel packing of Unc5D in the GPC3-mediated complex is reminiscent of the antiparallel arrangement in the Unc5D/Latrophilin3/FLRT2 complex, despite the otherwise different complex architectures and stoichiometries (Fig. S7E, F). This conformation may present a specific functional state of Unc5D, for example, it may cause distancing of the intracellular signalling domains of Unc5D molecules presented *in cis*, or allow an antiparallel *trans* interaction of Unc5D across different cells.

Post-translational glycan modifications such as C-mannosylation are emerging critical factors in receptor biology. For example, C-mannosylation and N-linked fucosylation are involved in mediating RTN4/NoGo interaction with the adhesion GPCR BAI in synapse formation and neuron-glia interaction (Wang et al., 2021). The interleukin-21 receptor is stabilised by an N-linked glycan that packs against mannosylated tryptophans within the same molecule (Hamming et al., 2012). The enzymatic C-mannosylation of Unc5 receptors by DPY19L1 is required for effective folding and stability of Unc5 TSP domains (Shcherbakova et al., 2017, 2019). We show here, for the first time, that C-mannosylated trytophans on the Unc5 TSP1 domain are involved in a novel and elaborate inter-chain glycan-glycan interaction, that takes place at the centre of the GPC3/Unc5 complex. Interestingly, Unc5 C-mannosylation was shown to play a role in neuroblast migration in *C.elegans* (Buettner et al., 2013). However, the site harbouring the hGPC3 N-linked glycosylation site that is required for the GPC3-interaction is conserved only in vertebrates and fly, not *C.elegans.* This suggests that the complex may not form in *C.elegans*, or assemble differently, perhaps involving Unc-40/DCC, which binds LON- 2/GPC3, and affects Unc5 signalling (Blanchette et al., 2015).

Nanobodies are increasingly used to modulate protein functions *in vitro* and *in vivo*, including also in clinical usage (Yang and Shah, 2020). Their small, compact, monovalent and rigid structure, and deep tissue penetration, make them ideal for use in a wide range of biological settings, often as targeting molecules. Here we show the effectiveness of Nano^glue^ and Nano^break^ in our biophysical binding assays and use them to modulate the function of Unc5-GPC3 *in vitro* and *in vivo*. We have also developed these nanobodies into tetrameric streptavidin-complexes, with enhanced binding capacities for functional assays, and as effective labelling agents *in situ* or in pull-down experiments. We expect these tools to be widely useful, given the broad expression patterns of GPC3 and Unc5, and the clinical relevance of both receptors (discussed below).

We have used two accessible *in vivo* cell migration systems to understand whether Unc5-GPC3 signalling impacts migrating cells. Neural crest-like neuroblastoma and cortical radial migration are established paradigms of cellular migration, with distinct characteristics: neuroblastoma cells undergo collective migration following a typical path through the embryonic tissue, while radial migration from the intermediate zone through the cortical plate relies on the interactions of individual neurons with their AP progenitor scaffolds. We found Unc5 and GPC3 are expressed in both systems: the neuroblastoma cells express both GPC3 and Unc5 receptors, with further Unc5 and GPC3 expressed in their environment. Young cortical neurons express mainly Unc5 receptors, while their AP scaffolds express GPC3. Unc5-GPC3 complex could be forming *in cis* or *in trans* in these systems (Fig. 7I and S7H, I). In the cortex, we find that enhancing or reducing GPC3-Unc5 interaction leads to impaired radial migration. This situation is reminiscent of previous studies showing that any alteration to the finely balanced adhesive or repulsive forces has a detrimental impact on the migration of these neurons (Seiradake et al., 2014; del Toro et al., 2020). It is possible that the reduction of Unc5-GPC3 interactions between APs and neurons removes a repulsive force that otherwise helps the neurons detach from their scaffold as they move forward, and thereby causes migration delays. Conversely, stabilizing the Unc5-GPC3 interaction artificially with Nano^glue^ could cause overly strong repulsion preventing the cells from attaching to the scaffold in the first place. Alternatively, we hypothesize that stabilizing the interaction interferes with the release mechanism from the GPC3-presenting scaffolds, which would be consistent with the complete lack of neural migration observed in the stripe assays, in the presence of Nano^glue^. Similar results have previously been obtained with other major guidance and adhesion systems, where cell migration speed can be reduced by modulating either adhesion or repulsion. Indeed, increased integrin-mediated adhesion to the extracellular matrix (Haage et al., 2020) or reducing Eph-EphrinB contact repulsion reduces cell motility (Rohani et al., 2011). Likewise, inhibiting fibronectin-integrin adhesion (Ramos and DeSimone, 1996) or increasing EphB-ephrinB repulsion (Wen and Winklbauer, 2017) impairs cell migration.

Unlike cortical neurons, many cancer cells display collective migration (te Boekhorst et al., 2016) (Piacentino et al., 2020). In our neuroblastoma model, GPC3-Unc5 signalling seems to act as a switch that determines cellular cohesion. Enhancing the interaction leads to more cohesive migration, whilst reducing it leads to the breaking up of the migrating cell cluster. Interestingly, cancer cells can reversibly switch from collective to individual migration mode, for optimal adaptation to their context (teBoekhorst et al., 2016, 2022). Modulation of GPC3-Unc5 interactions could thus contribute to mediating such opportunistic migration plasticity. The precise mechanism of neural crest cell targeting is still poorly understood, however our results suggest that GPC3-Unc5 interaction plays a key role in the targeting of neuroblastoma cells to the sympathoadrenal presumptive tissue. Reducing the interaction led to the premature stopping and forming of tumors within the migration path, while enhancing the interaction resulted in an overshoot of migration, with preferential tumorigenesis in the more distal adrenal part. Some tumors were found even further along this direction, migrating in a path normally taken by the enteric neural crest to target the developing gut. Taken together, and in analogy with the cortical migration paradigm, we find that GPC3-Unc5 signalling must be finely balanced to achieve effective collective migration and correct targeting, possibly because it could otherwise interfere with the perception of extracellular target recognition signals.

GPC3 and Unc5 are embedded within complex protein surface interactomes. For example, Unc5 receptors form complexes also with FLRTs (Seiradake et al., 2014) and Latrophilins (Jackson et al., 2016), which are known to contribute to cortical cell guidance. Given that these receptors are present in specific cortical cell populations (Seiradake et al., 2014; del Toro et al., 2020), there could be competition for the formation of different Unc5-signalling hubs. Also, cortical GPC3 could be interacting with receptors other than Unc5, as suggested by our *in vitro* stripe data. For example, GPC3 has been shown to bind Wnts (Capurro et al., 2005) and to promote the canonical Wnt/Beta-catenin pathway (Castillo et al., 2016), whose activation can provoke premature cortical migration (Woodhead et al., 2006). Wnt signaling also regulates neural crest migration, as does Latrophilin2 in different model organisms (Becker and Wilting, 2018; Yokote et al., 2019), and FLRT2 is expressed from early developmental stages in the trunk mesenchyme (Haines et al., 2006). Here, we have taken full advantage of the specificity of our structure-guided mutants, which retain most functional properties except for the protein binding-sites that were targeted, and our highly specific functional nanobodies. This approach has enabled us to focus our experiments on the novel Unc5-GPC3 interaction, despite the presence of other ligands. Many remaining questions can be answered using these tools, for example, regarding the roles of Unc5 and GPC3 in other tissues where these receptors are expressed, such as lungs, kidney (Iglesias et al., 2008; Liu et al., 2004; Schwab et al., 2003) and the vascular system (Freitas et al., 2008; Ng et al., 2009). Finally, the Unc5-GPC3-dependent mechanisms reported here during neuroblastoma migration could easily apply to other disseminating cancers, given that GPC3 is an oncofetal protein expressed by many pediatric solid embryonal tumors (Ortiz et al., 2019) and adult cancers (Li et al., 2018; Shimizu et al., 2019).

## Acknowledgements

We thank the Diamond Light Source and ESRF for beamtime, R. Klein for access to the proteomics facility of the Max Planck Institute of Biochemistry (Martinsried, Germany) and helpful discussion. Y. Shen for RNA data processing advice and M. Calvo from the Advanced Microscopy service (CCiT, university of Barcelona) for help with confocal microscopy. We thank Benjamin Villalard for assistance in the scRNAseq analysis. We thank the organisers of the CCP4-Diamond workshop during which some of the data presented here was collected. E.S. was funded by a Wellcome Trust Senior Research Fellowship (202827/Z/16/Z) and is supported by the EMBO Young Investigator Programme and the Oxford Kavli Institute for Nanoscience Discovery. VC was funded by the LabEx CORTEX and DEVWECAN of Université de Lyon, within the program “Investissements d’Avenir” (ANR-11-IDEX-0007) operated by the French National Research Agency (ANR) and the fondation Bettencourt-Schueller. DdT was funded by the Ramón y Cajal program (RYC-2017-23486) and MINECO project: RTI2018-095580-A-100. C.P. was funded by the FI fellowship from Generalitat de Cataluña and S.Z. was funded by the FPI fellowship from MINECO program. DC was funded by National Science Foundation (award 1755189), strategic research funds from the School of Biological Sciences at Victoria University of Wellington and RWJ Foundation grant 74260 to the Child Health Institute of New Jersey. Nanobody generation was funded by the John Fell Fund, University of Oxford and the Rosalind Franklin Institute EPSRC grant no. EP/S025243/1.

## Author contributions

OA led the crystallisation and initial crystallographic analysis of the proteins, designed mutants and produced samples for mass spectrometry and nanobody production.

CDB led the in vivo neuroblastoma work, the transwell migration assay, and RNA Seq analysis CP led the cortical in vivo work, pull downs, and stripe assays using cortical neurons MCO led stripe assays using cell lines, produced vectors and protein samples for *in vivo/in vitro* analysis, and contributed to SPR studies

MK led the characterisation of nanobodies and produced the streptavidin complexes MBS led cell aggregation and cell-based binding assays

MC led the MD simulation analysis

FR contributed to the *in ovo* neuroblastoma cell transplantations and the transwell migration assays

RR led the mass spectrometry analysis

JA led the advanced crystallographic data analysis and glycan refinement

MA oversaw the initial crystallographic data analysis and provided technical assistance EW contributed to the stripe assay analysis

EL oversaw crystallisation and crystallographic data collection DBA performed single cell RNA seq experiments

SZ performed the ISH experiments of Unc5-GPC3

PTNM modelled putative membrane protein complexes JH performed initial nanobody screening assays

DC and IP discovered the Unc5-GPC3 interaction in a protein interaction screen RO oversaw nanobody production

CVR oversaw mass spectrometry experiments

VC oversaw the work using neuroblastoma cells, and corresponding RNA seq analysis

DdT oversaw the work using primary neurons, cortical migration models, and corresponding RNA seq analysis

ES oversaw crystallographic, biophysical, nanobody and cell biology aspects of the work All authors have contributed to the manuscript.

## Methods

### Vectors and Cloning

We coned constructs of human GPC3 (cDNA clone BC035972) (hGPC3, residues 1-580; hGPC3^core^, residues 32-483, UG mutant, N241Q), mouse GPC3 (Uniprot ID: Q8CFZ4) (mGPC3^ecto^, residues 31-559; mGPC3^488^, residues 31-488; mGPC3^core^, residues 31-482), mouse Unc5A (Uniprot ID: Q8K1S4) (mUnc5A^ecto^, residues 1-359), mouse Unc5B (Uniprot ID: Q8K1S3) (mUnc5B, residues 26-934; mUnc5B^ecto^, residues 26-362), into the Age1-Kpn1 or EcoR1-Kpn1 cloning site of vectors from the pHLSec family (Aricescu et al., 2006). For protein purification, we used pHLSec vectors which also code for a C-terminal 6xHis-tag, for SPR we used a C-terminal Avi-tag. We used previously published rUnc5D constructs and derivatives thereof as indicated in the text (Jackson et al., 2015, 2016; Seiradake et al., 2014; del Toro et al., 2020), including (rUnc5D, rUnc5D^ecto^, rUnc5D^IgIgTSP^, rUnc5D^IgIg^, rUnc5D^TSPTSP^, FU mutant (W85N + S87T), human Unc5B (Q8IZJ1) (hUnc5B^ecto^), hUnc5A (Uniprot ID: Q6ZN44) (hUnc5A^ecto^, equivalent to Unc5A^Ig12T1^), mouse FLRT2 (FLRT2^LRR^). For cell binding and functional assays, full length constructs were used, either cloned into a pHLSec vector that encodes an intracellular mVenus, or the pCAGIG vector (Addgene), for visualisation. pCAGIG was modified to express mCherry instead of GFP, for certain experiments, and is then referred to as pCAGIC. For the nanobody expression and purification in WK6, Nano^glue^ and Nano^break^ in pADL-23c vector were used. Nanobodies were cloned into pHLsec C-terminal Avi-tag, for biotinylation in HEK293T cells. They were subcloned in the pCAGIG vector with the pHLSec-derived secretion signal, for *in vivo* experiments.

### Protein expression and purification

Recombinant protein expression and purification were performed as described (Aricescu et al., 2006; Seiradake et al., 2015). Briefly, adherent HEK293T or GnTI-deficient HEK293S cells were transiently transfected with the relevant plasmids using polyethylenimine (PEI) and grown for 5-10 days. Cell culture media were filtered to remove dead cells and buffer-exchanged to phosphate buffer saline (PBS) containing also 250 mM NaCl and 20 mM Tris (pH 7.5). Conditioned media were passed through HisTrap HP columns (GE Healthcare), washed with buffer supplemented with 40 mM imidazole and bound proteins were eluted using 20 mM Tris pH 7.5, 300 mM NaCl and 500 mM imidazole. The eluate was then subjected to size exclusion chromatography using Superdex 200 16/60 (GE Healthcare) in 10 mM Tris-HCl (pH 7.5) and 200 mM NaCl.

Nanobodies were expressed in E. coli strain WK6., grown at 37°C in Terrific Broth until an OD600=0.8. Expression was induced with 150 μM IPTG and incubation at 21°C for 16 h. Cells were harvested by centrifugation (6,000xg, 15 min). Cell pellets were resuspended and incubated in ice-cold 20% sucrose, 30 mM Tris-HCl, pH 7.5, 2 mM EDTA buffer for 20 min. The cell suspension was clarified by centrifugation (10.000 rpm, 20 min, 4°C). After the supernatant collection, cell pellets were resuspended and incubated in ice-cold 30 mM Tris- HCl, pH 7.5, 5 mM MgSO4 buffer for 20 min. The cell suspension was clarified by centrifugation (10.000 rpm, 20 min, 4°C). The supernatant was filtered, supplemented with 150 mM NaCl and 2 mM Imidazole and passed through HisTrap HP columns. The column was washed with 200 ml wash buffer (30 mM Tris-HCL, pH 7.5, 150 mM NaCl, 5 mM Imidazole) and the protein was eluted in elution buffer (30 mM Tris-HCL, pH 7.5, 150 mM NaCl, 500 mM Imidazole). Elution was loaded onto a Superdex200 16/600 HiLoad column in 20 mM Tris-HCL, pH 7.5, 200 mM NaCl.

For protein biotinylation in HEK293 cells, protein constructs in pHLsec C-terminal Avi-tag were co transfected with a vector encoding BirA (biotin ligase). Cell culture medium was supplemented with 100mM biotin. Proteins were purified as previously, but in ice cold conditions. For the nanobody-streptavidin complexes, streptavidin (Streptavidin-Alexa Fluor 594 conjugate, Thermo Fisher Scientific, or Streptavidin from Streptomyces avidinii, Sigma-Aldrich) was mixed with an excess of biotinylated nanobodies, incubated overnight at 4°C and subjected to size exclusion chromatography using Superdex 200 16/60 (GE Healthcare) in 20 mM Tris-HCl (pH 7.5) and 200 mM NaCl.

### Protein X-ray Crystallography

Proteins that were expressed in GnTI-deficient HEK293S cells were used for crystallisation trials. Crystals were grown by the vapor diffusion method at 18°C by mixing the protein solution and crystallization solution in a 1:1 ratio. Purified hGPC3^core^ was concentrated to 5.3 mg/ml, and crystals were obtained using crystallization solution 1 (20% ethylene glycol, 10% w/v PEG 8000, 0.1 M Tris/BICINE (pH 8.5) and 0.02 M of amino acids (L-Na-glutamate, alanine, glycine, lysine-HCl, and serine)). Crystals of mGPC3^core^ were obtained by concentrating the protein to 9.9 mg/ml in the presence of 100 mM NDSB256 and using crystallization solution 2 (0.2 M ammonium nitrate and 20% w/v PEG3350). Crystals of the complex hGPC3^core^ and rUnc5D^IgIgTSP^ were obtained by mixing the two proteins in a 1:1 molar ratio and concentrating them to 5.8 mg/ml. The protein solution was then mixed with crystallisation solution 1. Crystals of the mGPC3^488^ and rUnc5D^IgIgTSP^ complex were obtained by mixing the two proteins in a 1:1 molar ratio, concentrating to 7.1 mg/ml and mixing with 15% w/v PEG 3000, 20% v/v 1, 2, 4- butanetrol, 1% w/v NDSB 256, 0.1 M Gly-Gly/AMPD (pH 8.5) and 0.2 M of amino acids (DL-arginine HCl, DL-threonine, DL-histidine HCl H_2_O, DL-5-hydroxylysine HCl, trans-4-hydroxyl-L-proline).

### Structure determination

Crystals of hGPC3^core^, hGPC3^core^/rUnc5D^IgIgTSP^ and mGPC3^core^/rUnc5D^IgIgTSP^ were flash-cooled in their original crystallization condition. Crystals of mGPC3^core^ were cryoprotected by adding 20% v/v glycerol to the original crystallization solution. All diffraction data were collected at the Diamond Light Source synchrotron at 100K. Data was integrated using DIALS (via XIA2)(Winter et al., 2013, 2018), and integrated intensities were merged and scaled using programs from the CCP4 package (Winn et al., 2011). The data of hGPC3^core^/rUnc5D^IgIgTSP^ was also processed with Staraniso (Vonrhein et al., 2018). The structures were solved by molecular replacement (MR) using the models of h/mGPC3^core^, rUnc5D^IgIgTSP^ (Jackson et al., 2016) and PHASER (McCoy et al., 2007). Initial phases of mGPC3^core^ were obtained by MR using the central lobe of GPC1 and DLP structures (PDB 4YWT & 3ODN). We performed iterative cycles of model building and refinement in Phenix (Liebschner et al., 2019). Manual model building was performed in Coot (Emsley and Cowtan, 2004), and models were all atom refined using REFMAC (Murshudov et al., 2011) and Phenix (Liebschner et al., 2019). For the complexes, we used high-resolution models of individual components as targets, non-crystallographic symmetry (NCS), TLS and secondary structure restraints. The quality of the final models was assessed by MolProbity (Davis et al., 2007) and the CCP4i2 validation task (Potterton et al., 2018). Superpositions were done with SuperPose (Maiti et al., 2004).

### Glycan modelling, refinement and validation

N-glycans were built into positive omit electron density using the Coot N-linked carbohydrate building tool (Emsley and Crispin, 2018), then corrected manually where obvious discrepancies between map and model were encountered. The mannosylated tryptophans showed extra omit density consistent with this modification, which has been recently shown to force the mannoside moiety into a ^1^C_4_ conformation (an inverted chair) to keep the alpha linkage in a clash-avoiding equatorial conformation. In order to increase the observation to parameter ratio and restrain the mannoside’s ring conformation individually, external restraints for both N- and C-glycosylation were generated using the MKIV version of the Privateer software (Agirre et al., 2015). Restraints for the MAN-TRP covalent linkages were created using the AceDRG software (Long et al., 2017) through its CCP4i2 interface (Potterton et al., 2018). Glycans were iteratively refined using the REFMAC5 software and validated by Privateer software (Agirre et al., 2015).

### Molecular dynamics simulations and modelling

To simulate the hGPC3-rUnc5D complex, we followed essentially the same protocol as previously described to refine the structures of X-ray crystallography-derived complexes (Jackson et al., 2016; del Toro et al., 2020). Missing residues were modelled using MODELLER (Webb and Sali, 2016). As the purpose of this simulation was to identify protein-protein interactions, we removed the glycan parts. The proteins were solvated in TIP3P water with 150 mM NaCl. Molecular dynamics simulations were performed using GROMACS 2020 (Abraham et al., 2015) with the AMBER14SB force field (Maier et al., 2015). The system was first energy minimized and then equilibrated following a two steps procedure of constant temperature followed by constant pressure equilibrations (Lemkul, 2019). The 500 nanoseconds of production were run at 310 K and 1 bar in an NPT ensemble, using the velocity-rescaling thermostat (Bussi et al., 2007) coupled with the Parrinello–Rahman barostat (Parrinello and Rahman, 1981). To focus only on residue side chains movements, we keep the proteins backbone constrained while side chains were allowed to move freely during the course of the simulation. We used MDAnalysis (Michaud-Agrawal et al., 2011) to perform hydrogen bond analysis as previously described (del Toro et al., 2020). The Jupyter notebook used to perform such an analysis is available at: https://github.com/MChavent/Hbond-analysis. Briefly, we used a donor-acceptor distance cut-off of 3.0 Å and a cut-off angle of 120°. The hydrogen bond stability was defined as the percentage of the simulated time in which a residue forms stable hydrogen-bonds with its partner.

The models of membrane-bound GPC3 and Unc5D (Fig. S7H, I) were produced using structures presented here, and Alphafold models of the hGPC3 C-terminal region and the rUnc5D TSP2 and transmembrane domains (Jumper et al., 2021; Varadi et al., 2022). To position the Unc5D TSP2 domain, we ran 100 nanoseconds of atomistic simulations on the Alpha-fold model containing Ig2+TSP1+TSP2 (as described above) without constraints. While the Ig1 -TSP1 linkage was stable in these simulations, we observed that the TSP2 domain explored a range of positions relative to TSP1, possibly due to the presence of a proline residue in the linker between TSP1 and TSP2. We extracted 7 representative structures from the molecular dynamics trajectory, shown superposed in Fig. S7G. We used the MultiSeq VMD plugin (Roberts et al., 2006) to superimpose via the TSP1 domains for this figure. Missing linkers for Unc5D and GPC3 were added using MODELLER (Webb and Sali, 2016). We added heparan sulphate glycans and GPI anchors using the CHARMM-GUI glycan modeller (Park et al., 2019).

### SPR

Equilibrium binding experiments were performed at 25°C using a Biacore T200 instrument (GE Healthcare) using PBS + 0.005% (v/v) polysorbate 20 (pH 7.5) or 20mM Tris + 200mM NaCl + 0.005% (v/v) polysorbate 20 (pH 7.5) as running buffers. The regeneration buffer was 2 M MgCl_2_. Glypican, Unc5 or nanobody constructs were biotinylated enzymatically at a C-terminal Avi-Tag and coupled to a streptavidin-coated CM5 chip. Data were analysed using the BIAevaluation software. Indicative K_D_ and R_max_ values were obtained by nonlinear curve fitting of a 1:1 Langmuir interaction model (bound = R_max_/(K_D_ +C), where C is analyte concentration calculated as monomer.

### Cell Binding Assay

HEK293T cells were transfected using mVenus-tagged constructs. Eighteen hours after transfections, cells were incubated with buffer (HBSS with 1% BSA and 10 mM HEPES (pH 7.5)) for 30 minutes on ice, and then with buffer containing 250 nM purified His-tagged protein that was previously pre-clustered with anti-His (mouse; Thermo Fisher Scientific) in a 2:1 (protein:antibody) ratio for 60 minutes on ice. Cells were then washed with PBS and fixed with 4% PFA. Cells were then incubated with anti-mouse-Cy3 in buffer for 60 minutes on ice. The cells were washed with PBS, stained with DAPI and mounted using Immu-Mount. Analysis was performed in ImageJ (Schneider et al., 2012).

### Native Mass Spectrometry Experiments

Unliganded mGPC3^core^ and rUnc5D^IgIgTSP^ were concentrated separately to 10 µM, dialysed against 1M ammonium acetate buffer (pH 7.5) overnight at 4 °C, and injected at 3 µM concentration. The mGPC3^core(R355A/R358A)^ - rUnc5D^IgIgTSP^ complex was concentrated to 15 µM (assuming a 4:4 stoichiometry) and dialysed against 200mM ammonium acetate (pH 7.5) and injected at 3.3 µM concentration. The protein samples were loaded into in-house prepared gold-coated capillary needles (Harvard Apparatus) and were injected directly to the mass spectrometer. The experiments were performed using a Q-Exactive UHMR Hybrid Quadrupole-Orbitrap mass spectrometer (Thermo Fisher). Typically, 3 μl of protein solution was electrosprayed from gold-coated capillaries. The instrument parameters for MS are as follows: 1.2 kV capillary voltage, S-lens RF 200%, quadrupole selection from 1,500 to 20,000 *m*/*z* range, in-source trapping energy (0-20V), nitrogen UHV pressure of 6.07 × 10^−10^ mbar and capillary temperature of 100 °C. The resolution of the instrument was 17,500 at *m*/*z* = 200 (transient time of 64 ms). The noise level was set at 3 rather than the default value of 4.64. For MS/MS analysis, collisional activation in the HCD cell was provided (0-300 V). Calibration of the instruments was performed using a 10 mg/ml solution of cesium iodide in water. Data were analyzed using the Xcalibur 4.1 (Thermo Scientific).

### LC-MS/MS Identification of Tryptophan Mannosylation

Tryptophan mannosylation was verified by LC-MS/MS. 5 μg of each protein (1 mg/ml) was diluted to a final volume of 100 μL in denaturing buffer (8 M Urea, 50 mM Ammonium Bicarbonate). The sample was reduced with DTT (2 μL of 200 mM solution in denaturing buffer) at 56 °C for 25 minutes, followed by alkylation with iodoacetamide (4 μL of 200 mM solution in denaturing buffer) at room temperature for 30 minutes. Alkylation was quenched by further addition of 2 μL of DTT solution. The samples were further diluted three-folds using 50 mM Ammonium Acetate buffer. Trypsin was added to the sample in 1:50 (Enzyme: Protein (w/w)) ratio. The sample was incubated at 37 °C for 16 hours. Next day, the digested sample was divided into two fractions. One of the fraction was quenched using 10% Formic acid solution. To the other fraction, AspN was added in 1:20 (Enzyme: Protein (w/w)) ratio. The sample was incubated further at 37 °C for 4 hours after which it was quenched using 10 % Formic acid solution. Resulting peptides were analyzed on an UltiMate 3000 UHPLC system (Thermo Fisher) connected to an Orbitrap Eclipse Tribrid mass spectrometer (Thermo Fisher). The peptides were trapped on an guard column (Acclaim PepMap 100, 75 µm x 2 cm, nano viper, C18, 3 µm, 100 Å, Thermo Fisher) using solvent A (0.1% Formic acid, water). The peptides were separated on an Acclaim PepMap analytical column (75 µm × 150 mm, RSLC C18, 3 µm, 100 Å) using a linear gradient (length: 90 minutes, 6 % to 45 % solvent B (0.1% formic acid, 80 % acetonitrile, 20% water), flow rate: 300 nL/min). The separated peptides were electrosprayed directly into the mass spectrometer in the positive ion mode using data-dependent acquisition with a 3 second cycle time. Precursors and products were detected in the Orbitrap analyzer at a resolving power of 60,000 and 30,000 (@ m/z 200), respectively. Precursor signals with an intensity >1.0 x 10^-4^ and charge state between 2-7 were isolated with the quadrupole using a 0.7 m/z isolation window (0.5 m/z offset) and subjected to MS/MS fragmentation using higher-energy collision induced dissociation (30% relative fragmentation energy). MS/MS scans were collected at an AGC setting of 1.0 x 10^4^ or a maximum fill time of 100 ms and precursors within 10 ppm were dynamically excluded for 30 seconds.

Raw data files were processed using MaxQuant software (Version 1.6.3.4), having in-built Andromeda search engine (Cox and Mann, 2008; Cox et al., 2011). The peak lists were searched against individual Unc and common contaminant proteins. Carbamidomethylation was kept as fixed modification whereas acetylation (protein N-term), oxidation (methionine), and hexose (tryptophan) were used as variable modifications. Protein and peptide false discovery rate was kept at 1%. Trypsin and AspN were set as the protease and up to four missed cleavages were allowed.

### Nanobody Generation

Antibodies to hGPC3^core^ were raised in a llama by intra-muscular immunization with purified protein using Gerbu LQ#3000 as the adjuvant. Immunisations and handling of the llama were performed under the authority of the project license PA1FB163A (University of Reading, UK). Total RNA was extracted from peripheral blood mononuclear cells, and VHH complementary DNAs were generated by RT-PCR. The pool of VHH-encoding sequences was amplified by two rounds of nested PCR and cloned into the SfiI sites of the phagemid vector pADL-23c as previously described (Huo et al., 2021). Electrocompetent E. coli TG1 cells were transformed with the recombinant pADL-23c vector, and the resulting TG1 library stock was infected with M13K07 helper phage to obtain a library of VHH-presenting phages. Phages displaying VHHs specific for hGPC3^core^ were enriched via two rounds of bio-panning on biotinylated hGPC3^core^, and individual phagemid clones were picked. VHH-displaying phages were recovered by infection with M13K07 helper phage and tested for binding to hGPC3^core^ by enzyme-linked immunosorbent assay (ELISA). Phage binders were ranked according to the ELISA signal and grouped by CDR3 sequence identity.

### Nanobody expression and purification

Nanobodies were expressed in E. coli strain WK6., grown at 37°C in Terrific Broth until an OD600=0.8. Expression was induced with 150 μM IPTG and incubation at 21°C for 16 h. Cells were harvested by centrifugation (6,000xg, 15 min). Cell pellets were resuspended and incubated in ice-cold 20% sucrose, 30 mM Tris-HCl, pH 7.5, 2 mM EDTA buffer for 20 min. The cell suspension was clarified by centrifugation (10.000 rpm, 20 min, 4°C). After the supernatant collection, cell pellets were resuspended and incubated in ice-cold 30 mM Tris- HCl, pH 7.5, 5 mM MgSO4 buffer for 20 min. The cell suspension was clarified by centrifugation (10.000 rpm, 20 min, 4°C). The supernatant was filtered, supplemented with 150 mM NaCl and 2 mM Imidazole and passed through HisTrap HP columns. The column was washed with 200 ml wash buffer (30 mM Tris-HCL, pH 7.5, 150 mM NaCl, 5 mM Imidazole) and the protein was eluted in elution buffer (30 mM Tris-HCL, pH 7.5, 150 mM NaCl, 500 mM Imidazole). Elution was loaded onto a Superdex200 16/600 HiLoad column in 20 mM Tris-HCL, pH 7.5, 200 mM NaCl.

### Cell aggregation assay

K562 suspension cells were cultured in RPMI-1640 media supplemented with 10% FBS and 5% L-Glutamine. Cells at a concentration of 2x10^7^ cells/ml were transfected with control pCAGIG/gCAGIC plasmids, or those coding for Unc5 or GPC3 constructs using the Neon transfection system for electroporation (Settings: 1450V, 3 pulses, 10 ms). Twenty-four hours after transfection, cells were harvested, passed through a 40 µm cell-strainer and used at a concentration of either 2 x 10^5^ cells/ml or 4 x 10^5^ cells/ml in aggregation media (Neurobasal-A media supplemented with 2 mM L-glutamine, 10% FBS, 4% B-27 and 20 mM HEPES). For the competition experiments, different amounts of nanobodies (5, 10 and 50 µg) were added at this stage. Cells were then left to aggregate at 37°C, 5% CO_2_ and 250 rpm for 90 minutes. After the incubation, cells were diluted in 2 ml of PBS and imaged in a 6-well plate using a Nikon ECLIPSE TE2000-U inverted fluorescence microscope. Images presented were obtained using Inverted DeltaVision widefield microscope at 37°C. The total area of cells and the total area of the aggregates for each picture were calculated using the Analyze particle tool in ImageJ. The threshold used to distinguish cells and aggregates was determined at 1000 pixel^2^ (>3 cells).

### K562 protein expression tests

For surface staining, K562 cells were harvested 24 hours after electroporation and cooled to 4°C. The cells were then incubated with blocking buffer: HBSS with 1% BSA and 10 mM Hepes (pH 7.5) for 30 minutes. For Unc5D expressing cells, cells were incubated with anti-HA (mouse; Sigma-Aldrich) antibody that was pre-clustered with secondary antibody-Cy5 for 40 minutes (ratio 1:7.5 for primary:secondary antibody). For GPC3 expressing cells, cells were incubated anti-FLAG (rabbit; Sigma-Aldrich) antibody that was pre-clustered with secondary antibody conjugated with Alexa 647 for 40 minutes (1:7.5 ratio). Cells were washed with PBS, fixed with 4% PFA, DAPI-stained, washed and resuspended in 30% sucrose/PBS. Cells were then deposited onto microscope slides using a homemade cytospin (Sisino et al., 2006). Imaging was performed using an Inverted DeltaVision widefield microscope with CCD. Colocalization for the surface quantification was performed using ImageJ (Schneider et al., 2012). For the total amount of protein, K562 cells expressing the desired constructs were harvested 24 hours after electroporation, lysed by sonication, and analysed on SDS-page/western blot with mouse anti-His, rabbit anti-FLAG, mouse anti-HA or mouse anti-Actin.

### Analysis of published single cell RNASeq dataset

Single Cell RNASeq of human neuroblastoma tumor samples were exploited from Dong et al. (2020; GEO ID: GSE137804) public dataset. Raw sequencing data were processed following the partially published method details, explaining the different UMAP obtained in the present study. The R package Seurat (v4.0.1) was used to calculate the quality control metrics. To filter out low quality cells, we kept all cells with at least 200 detected genes and less than 10% of mitochondrial genes. Doublet cells were removed with the R package DoubletFinder (v2.0.3). To merge all samples without biasing the analysis with batch effects, while preserving the biological variation, we applied Seurat integration and re-computed a clustering based on the corrected matrix. Single RNAseq data for cortex samples were obtained from the published NCBI Gene Expression Omnibus with accession numbers GSE65000 (Florio et al., 2015)and GSE153164 (di Bella et al., 2021). We used the same UMAP coordinates and metadata information with the cluster categorization provided by the authors.

### Chick embryos

Naked Neck strain embryonated eggs were obtained from a local supplier (Elevage avicole du Grand Buisson, Saint Maurice sur Dargoire, France). Laying hen’s sanitary status was regularly checked by the supplier according to French laws. Eggs were housed at 18°C until use. They were then incubated at 38.5°C in a humidified incubator until the desired developmental stage, i.e, HH14 for the graft step (54 hours of incubation).

### Cell lines used for grafting

SY5Y and SY5Y:GFP (Delloye-Bourgeois et al., 2017) NB cell lines were cultured in Dulbecco’s Modified Eagle Medium (DMEM) GlutaMAX™ (Life Technologies). Media were each supplemented with 10% Fetal Bovine Serum (FBS), 25 U/mL Penicillin Streptomycin (Gibco), 2.5 µg/mL Amphotericin B (Sigma-Aldrich). Cell lines were maintained in sterile conditions in a 37°C, 5% CO2-incubator. Cell lines have not been re-authenticated for the present paper.

### Plasmids, siRNAs, cell transfection for grafting

Control siRNA (siRNA scr) (siRNA Universal Negative Control #1 SIC001) and human GPC3 siRNA (NM_004484; SASI_Hs01_00205845) were purchased from Sigma-Aldrich and used at a concentration of 50 nM. Vectors encoding for nanobodies were used at a concentration of 2 µg/mL. For siRNA and plasmids transfection, cells were transfected with JetPrime according to the manufacturer’s guidelines (PolyPlus).

### RNA Isolation and Quantitative Real-Time PCR (qRT-PCR)

For qRT-PCR analysis, total RNA was extracted from cells using the Nucleospin RNAII kit (Macherey-Nagel). One µg of total RNA was reverse-transcribed using the iScript cDNA Synthesis Kit (BioRad). qRT-PCR was performed using the LightCycler480 SYBRGreen I Master1 kit (Roche Life Science) and the CFX Connect Real-Time PCR Detection System (BioRad). The following list of primers was used in the study:

Human HPRT: Fwd: for TGACACTGGCAAAACAATGCA/ Rev: GGTCCTTTTCACCAGCAAGCT;

Human UNC5A: PrimerPCR SYBR Green Assay; qHsaCID0013056

Human UNC5B: PrimerPCR SYBR Green Assay; qHsaCID0021074

Human UNC5C: PrimerPCR SYBR Green Assay; qHsaCID0016268

Human UNC5D: PrimerPCR SYBR Green Assay; qHsaCED0045738

Human GPC3: PrimerPCR SYBR Green Assay; qHsaCID0016381

### Immunofluorescence on chick embryo slices

Chick embryos of interest were harvested and fixed in 4% paraformaldehyde (PFA). Embryos were embedded in 7.5% gelatin and 15% sucrose in PBS to perform 20 µm transverse cryosections. Permeabilization and saturation of sections were performed in PBS with 3% BSA and 0.5% Triton. The following primary antibodies were applied to sections: anti-HNK1 mouse IgM (1/50, 3H5, DSHB), anti-GFP rabbit IgG (1/500, Thermo Fisher Scientific, #A11222), anti-mitochondria mouse IgG (1/500, Millipore #MAB1273) and the following secondary antibodies: Alexa 647 anti-mouse IgM (1/500, Thermo Fisher Scientific #A21238); Alexa 488 anti-rabbit IgG (1/500, Thermo Fisher Scientific, #A11008), Alexa 555 anti-mouse IgG (1/500, Thermo Fisher Scientific #A31570) and. Nuclei were stained with Hoechst (Thermo Fisher Scientific, H21486). Slices were imaged with a confocal microscope (Olympus, FV1000, X81) using a 10X objective. The position of isolated cells and tumor masses was pointed on a reference image for each slice in which tumor cells could be detected, using Fiji software.

### Transwell migration assays

Transfected SY5Y cells were plated on the porous filter of the upper chamber of transwell culture dishes (8 μm pore size; BD Falcon, NJ, 5 x 10^4^ cells/ml). The cells were then incubated for 60 hours in a 37°C, 5% CO2-incubator. Cells retained on the upper face of the membrane were scrubbed using cotton swabs. The transwell culture dishes were then fixed with with 4% PFA for 30 minutes, before washing with 3 successive PBS-baths and mounting in Mowiol. Migrating cells were counted using a confocal microscope (Olympus, FV1000, X81).

### Stripe assays

We prepared the stripe assays essentially as previously described (del Toro et al., 2020). 50 μg/ml of Fc recombinant protein, GPC3^core^ or GPC3^coreUG^, were mixed with Alexa594- conjugated anti-hFc antibody (Thermo Fisher Scientific, A11014) in PBS. Proteins were injected into matrices (90 μm width) (17546017) and placed on 60 mm dishes, resulting in red fluorescent stripes. After 30 min incubation at 37°C, dishes were washed with PBS and matrices removed. Dishes were coated with 50 μg/ml Fc or GPC3^coreUG^ protein mixed with 120 μg/ml anti-hFc (Jackson ImmunoResearch 109-005-098) for 30 min at 37°C and washed with PBS. Stripes were further coated with 20 μg/ml Laminin in PBS for at least 2 hours and washed with PBS. Cortical neurons (E15.5) were cultured on the stripes in Neurobasal medium supplemented with B27 (Gibco). For testing the effects of nanobodies, neurons were cultured in medium containing 50 μg/ml streptavidin Alexa 594 (CN), Strep-Nano^break^ or Strep-Nano^glue^. After 24 hours neurons were fixed with 4% PFA in PBS for 20 min at room temperature (RT). Neurons were washed and incubated with rabbit monoclonal anti-beta-III tubulin antibody (Sigma-Aldrich, T2200) after 20 min permeabilization in 1% BSA, 0.1% Triton X-100 in PBS. Cy2 anti-rabbit IgG secondary antibody (Jackson ImmunoResearch, cat#111-225-144) was used to visualize the tubulin signal. Nuclei were counterstained with DAPI before mounting. The numbers of beta-III-tubulin-positive (green) pixels on red or black stripes were quantified with ImageJ (version 1.53f) (Schneider et al., 2012) using a custom-made automatic macro that is available upon request.

For cell lines, 50 μg/ml of GPC3^core^ protein was mixed with Alexa568-conjugated anti-hIgG antibody (Thermo Fisher Scientific, A21090) in PBS. Protein was injected into matrices (90 μm width) (17546017) and placed on 60 mm dishes, resulting in red fluorescent stripes. After 30 min incubation at 37°C, dishes were washed with PBS and matrices removed. Dishes were coated with 50 μg/ml GPC3^coreUG^ protein mixed with 120 μg/ml anti-hFc (Jackson ImmunoResearch 109-005-098) for 30 min at 37°C and washed with PBS. HeLa, SY5Y or N2A cells were cultured on the stripes in 2% FBS medium, with 50 μg/ml Strep-Nano^break^ or Strep- Nano^glue^, or without nanobody (control). After 16 hours cells were fixed with 4% PFA in PBS for 20 min at room temperature and nuclei were counterstained with DAPI before mounting and imaging. The number of DAPI positive pixels on red or black stripes was quantified.

### RNA In Situ Hybridization (ISH) and Immunohistochemistry

Embryonic brains were fixed in 4% PFA over-night. 10 μm Cryo-sections were pre-treated using the RNAscope Universal Pretreatment Kit (Advanced Cell Diagnostics, Cat#322380). RNA In Situ Hybridizations (ISH) were performed using the RNAscope Fluorescent Multiplex Reagent Kit (Advanced Cell Diagnostics, Cat#320850) according to manufacturer’s instructions. The target genes (Unc5D and GPC3) are listed in the Key Resources Table. Following ISH, sections were immunostained using mouse anti-Pvim 1/300 (Abcam, Cat#ab20346) and rat anti-Ctip2 1/600 (Abcam, Cat#ab123449) in combination with the Alexa Fluor 488 secondary antibodies 1/200. Images were acquired using a Zeiss LSM880 confocal laser scanning microscope and processed with ImageJ software.

### In utero electroporation

In utero electroporation was performed at E13.5 with anesthetized C57BL/6 mice as previously described (del Toro et al., 2020). DNA plasmids were used at 2 µg/µl and mixed with 1% fast green (Sigma-Aldrich, final concentration 0.2%). Plasmids were injected into the ventricle with a pump-controlled micropipette. After injection, six 50 ms electric pulses were generated with electrodes confronting the uterus above the ventricle. The abdominal wall and skin were sewed, and the mice were kept until E16.5 embryonic stage. To knockdown GPC3 by shRNA *in vivo*, we used the best out of 3 different tested target regions embedded in the in the pCAG-miR30 vector, with the following sequence: GCCGAAGAAGGGAACTGATTC. This shRNA was validated in HEK293T cells, by co-transfection with GPC3 followed by western blotting. Secreted versions of Unc5D extracellular domains (Unc5D^IgIgTSP^ and Unc5D^IgIgTSPGU^) as well as the nanobodies (Nano^glue^ and Nano^break^) were cloned into the pCAGIG vector. The expression of all constructs was validated by expression in HEK293T cells and analyzed on western blots.

### Pull-down experiments

Pull-down experiments were performed as previously described (del Toro et al., 2020). Fresh E15.5 mouse cortices were homogenized for 1 min at 4°C with an electric homogenizer using the following lysis buffer: 50 mM Tris-HCl (pH 7.4), 150mM NaCl, 2mM EDTA, 1% Triton X-100 and protease inhibitors (Sigma-Aldrich 04693132001). Samples were incubated on ice for 20 min and centrifuged for 10 min at 3000 rpm. Supernatant was collected and protein was measured using the Bio-Rad protein assay (Biorad, 5000001). 1.5 mg of protein at a final concentration of 2 μg/μl in lysis buffer (volume: 750 μl) was used for each pull-down. Control pull-down contained lysate and 40ul of high-capacity streptavidin agarose resin (Thermo Fisher Scientific, 20357, 50% w/v), whereas the nanobody condition contained the same beads plus 2μg of biotinylated Nano^glue^. Samples were incubated overnight at 4°C under rotatory agitation. The next day, agarose beads were centrifuged for 5 min at 3000 rpm and washed three times (first wash with 400 μl of lysis buffer, second wash with 1:1 (v/v) lysis buffer:PBS, last wash only PBS). Pulled-down samples were processed for mass spectrometry (MaxQuant run, Proteomic facility, Max Planck Institute of Biochemistry, Martinsried, Germany). Volcano plots were generated using the DEP package in R-studio.

Pull-down experiments to investigate the effects of Nano^glue^ and Nano^break^ on the Unc5 – GPC3 interaction were performed by coupling biotinylated hGPC3^core^ to high-capacity streptavidin agarose resin (Thermo Fisher Scientific, 20357). Unc5^ecto^ was mixed with nanobody in 1:1, 1:5, and 1:10 molar ratios in 20mM Tris (pH7.5), 200mM NaCl, 1% BSA and incubated with hGPC3^core^-coated strep beads. Beads were washed with 20mM Tris (pH7.5), 200mM NaCl and the proteins were eluted with SDS-containing gel-loading buffer + 5% beta-mercaptoethanol. Samples were analysed using SDS-PAGE.

---

**Fig. S1.**
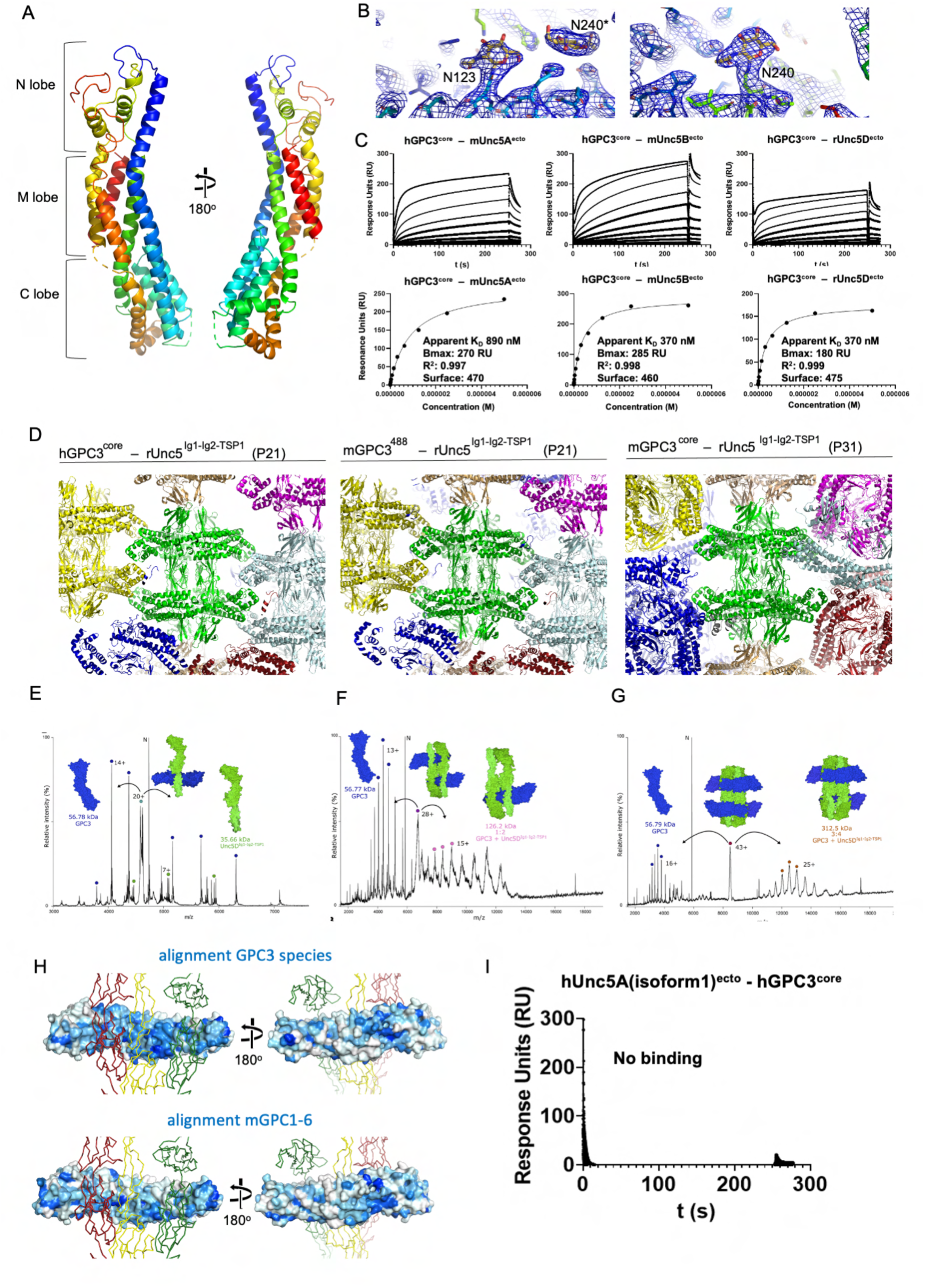
mGPC3^core^ structure and GPC3-Unc5D complex data. **(A)** Mouse GPC3^core^ structure coloured according to the rainbow (blue: N-terminus, red: C-terminus). **(B)** Electron density map calculated from murine GPC3^core^ crystals is shown in blue, centered on N123 and N240 of a symmetry-related molecule* (left) and N240 (right). **(C)** SPR experiments show binding of Unc5 extracellular domains to hGPC3^core^. The apparent K_D_s (K_Dcalc_) were calculated using a 1:1 binding model and are indicative only. Bmax, R^2^ and amount of ligand immobilised on the flowcell surface are indicated. **(D)** Crystal packing environment for the three complex structures. Each octameric unit is shown in a different colour, with a central unit in green. **(E-G)** Tandem MS (MS/MS) analysis of peaks presented in Fig. 2A. Peaks reveal rUnc5^IgIgTSP^ and hGPC3^core(R355A/R358A)^ subcomplexes. The 93 kDa peak dissociated into masses corresponding to GPC3 (56.78 kDa excluding glycans) and Unc5D (35.66 kDa excluding glycans), the peaks corresponding to 185 and 370 kDa dissociated into GPC3 (56.78 kDa) and a mass of 126 kDa (consistent with a 2:1 Unc5D:GPC3 complex). In the 370 kDa peak we additionally detected a 312 kDa species (consistent with a 4:3 Unc5D:GPC3 complex). **(H)** rUnc5D^IgIgTSP^ is shown in red, yellow and green ribbons, as found in the complex with mGPC3^core^. The surface of mGPC3^core^ is coloured in shades of blue according to sequence conservation (blue = conserved, while = not conserved). Surface conservation was calculated using aligned sequences from human, mouse, opossum, chicken, frog, and fish GPC3 (top) or mouse GPC1-6 (bottom). Note that the Unc5-binding site is less conserved amongst mouse GPC1-6 sequences, compared to different GPC3 sequences. **(I)** SPR results show that hUnc5A isoform A, which lacks a TSP1 domain, is unable to bind hGPC3^core^.

**Fig. S2.**
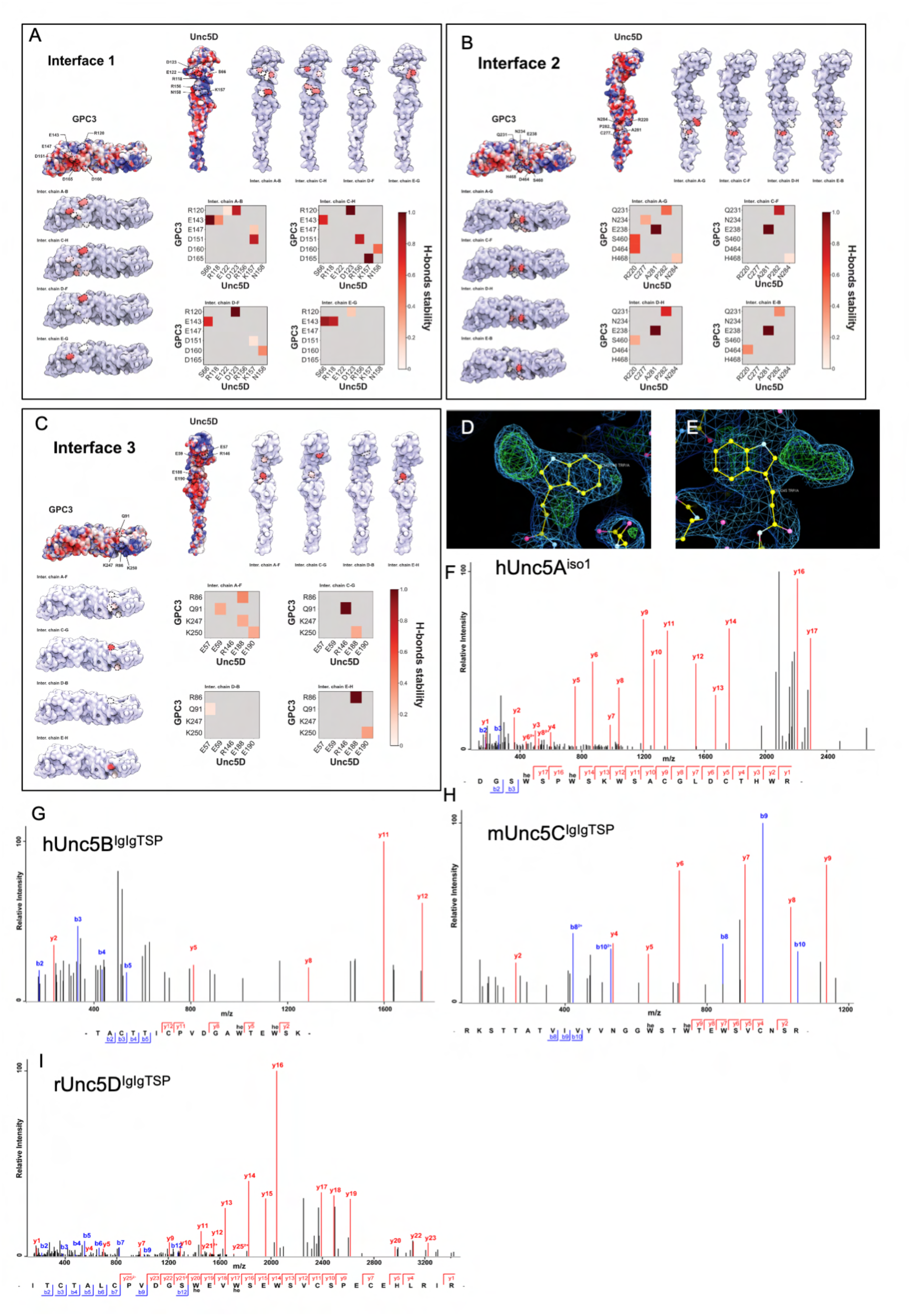
Hydrogen bond analysis during MD simulation and mass spectrometry analysis of Unc5 peptides. **(A-C)** Quantification of MD simulation results for each of the four pseudo-symmetrical copies in the complex, for each of the three rUnc5D- hGPC3 interfaces described in Fig. 2. **(D, E)** Views of the electron density maps calculated for the X-ray crystal structure of Unc5A^iso1^ (Seiradake et al. Neuron 2014): the 2Fo-Fc is shown in blue (1 sigma level). The Fo-Fc map is shown in red/green (+/- 3 sigma level). Extra density is observed on the first two of the TSP tryptophans of the consensus W_1_xxW_2_xxW_3_ motif. **(F-J)** LCMSMS of the tryptic Unc5 peptides confirming the C-mannosylation of tryptophan residues in the TSP1 domains of Unc5 proteins expressed in HEK cells.

**Fig. S3.**
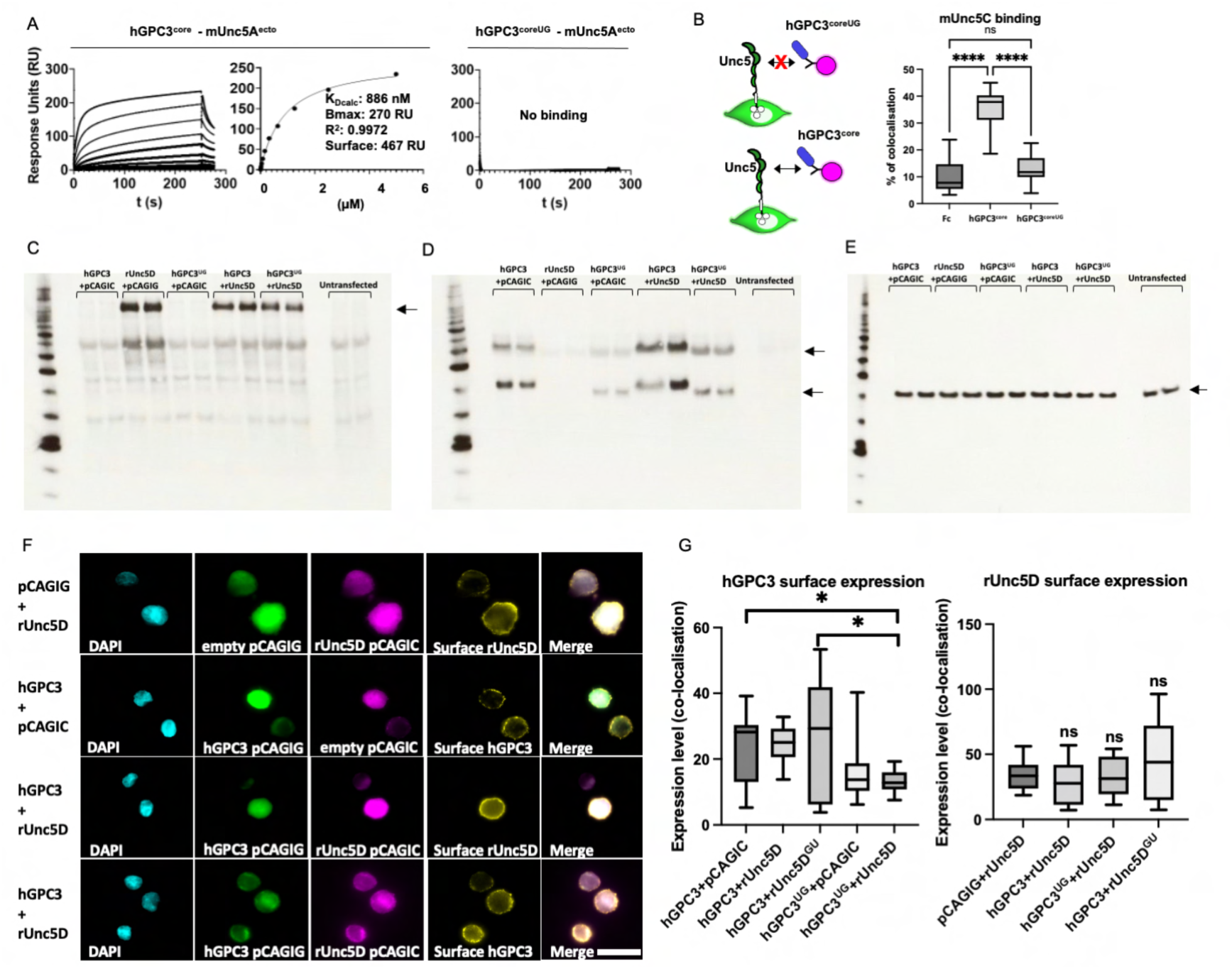
mUnc5A and mUnc5C binding results, protein co-expression analysis. **(A)** SPR results show binding of hGPC3^core^ protein to mouse Unc5A ectodomain. The apparent K_D_ (K_Dcalc_) for the wild type protein interaction was calculated using a 1:1 binding model and is indicative only. Bmax, R^2^ and the units of ligand immobilised on the flowcell surface are indicated. The N241Q mutant protein (hGPC3^coreUG^) does not show binding. **(B)** We used a cell-based assay to show that hGPC3^core^, but not the mutant, binds to mUnc5C expressed on cells. **(C)** Western blot analysis using anti-HA to visualise HA-tagged Unc5 constructs expressed in cell aggregation assays. **(D)** Same samples as in panel C, but here visualising Flag-GPC3. **(E)** Same as panel C, but using anti-actin control. **(F)** Cell-surface staining using anti-HA and anti-Flag was performed to complement the total protein expression analysis shown in panels C-E, and to include additional conditions. Representative images are shown. Scale bars = 30 µm. **(G)** Quantification of the experiments shown in panel F.

**Fig. S4.**
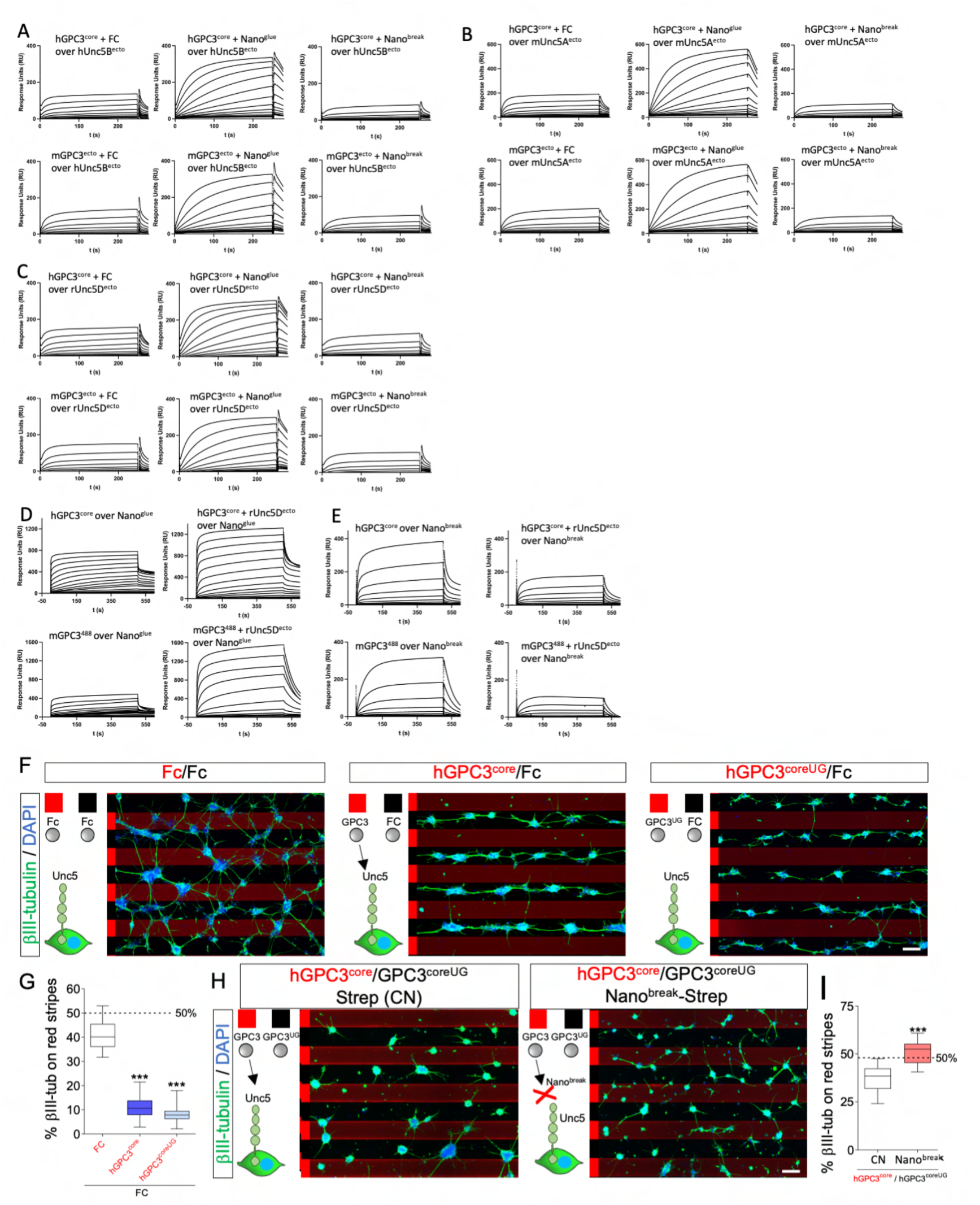
Nano^glue^ and Nano^break^ in SPR experiments and stripe assays. **(A-C)** Binding curves from SPR experiments. Unc5A,B or D receptor ectodomains were immobilised. Human or murine GPC3^core^ was injected using a 2-fold dilution series (top concentrations are 4.5 μM), in the presence or absence of Nano^glue^ or Nano^break^. The concentration of the nanobody was kept constant (9 μM when mixed with hGPC3^core^, 4.5 μM when mixed with mGPC3^ecto^). **(D, E)** An analogous experiment was performed using immobilised nanobodies, and different concentrations of human or murine GPC3^core^ and Unc5D^ecto^. Taken together, the results demonstrate that Nano^break^ competes with Unc5 for GPC3-binding, whilst Nano^glue^ strengthens the interaction. Calculated K_D_s for nanobody-GPC3 interactions are shown in Fig. 4C. Given the unusual stoichiometry of the Unc5-GPC3 complex, we have not calculated K_D_ values from experiments containing also Unc5. **(F)** Purified proteins were immobilised in a stripe pattern to assess their effect on the migration of cortical neurons. GPC3^core^ and GPC3^coreUG^ trigger strong cell repulsion, compared to neutral control protein (Fc). **(G)** Quantification of the experiments shown in panel F. **(H)** We performed GPC3^core^/GPC3^coreUG^ stripe assays, but in the presence of streptavidin (CN) or streptavidin-nanobody complexes. Nano^break^ reduced the ability of neurons to distinguish between hGCP3^core^ and hGCP3^coreUG^. **(I)** Quantification of data shown in panel H. ***p < 0.001, two-tailed Student’s T test. Scale bar represents 90 μm (F,H).

**Fig. S5.**
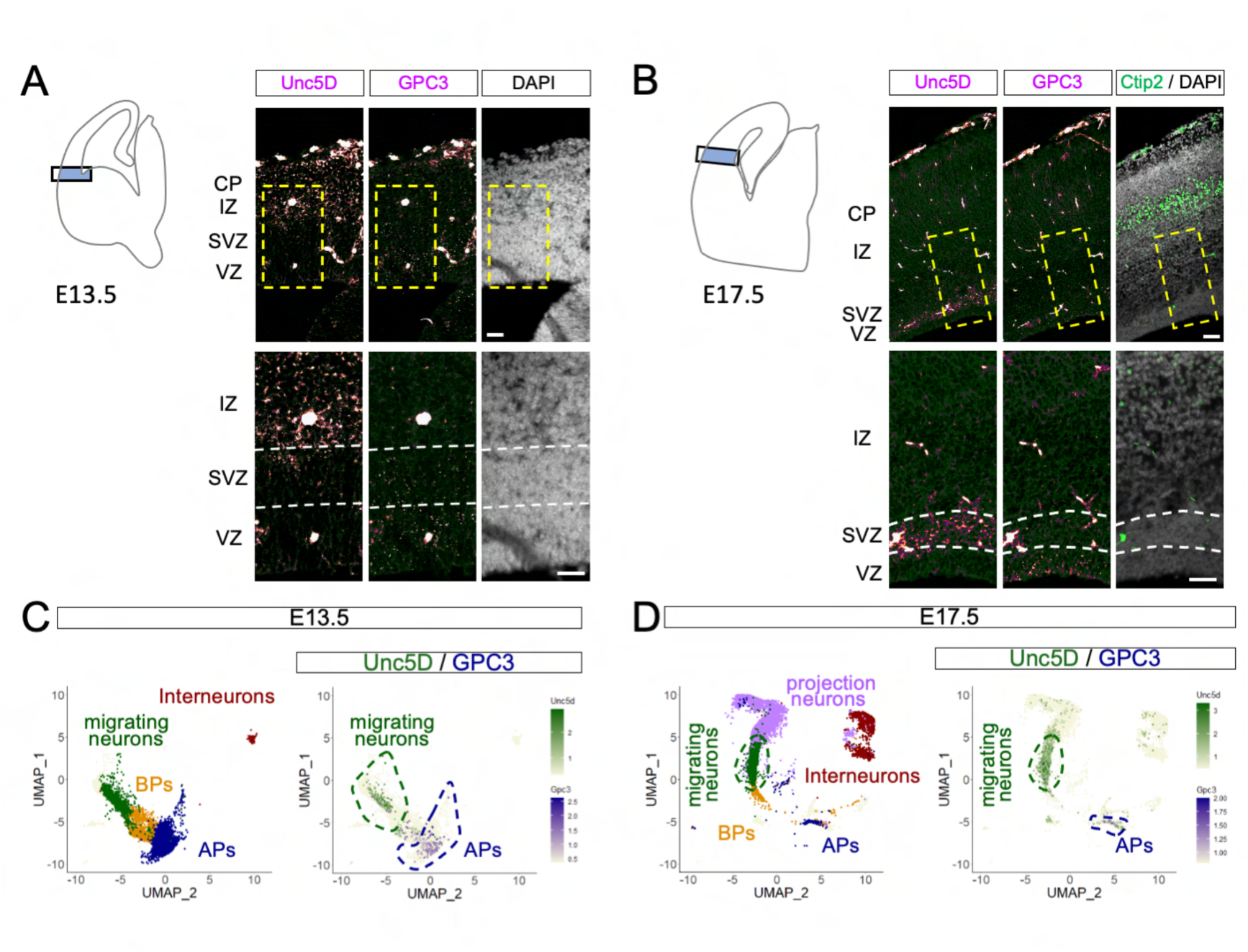
Unc5D and GPC3 are expressed during cortical development. **(A-B)** ISH for Unc5D and GPC3, colored in magenta, shows expression in the cortex of coronal sections of E13.5 (A) and E17.5 (B) mouse embryos. Each panel shows a diagram on the left, which is indicating the cortical region shown. The area in the dashed rectangle is magnified on the bottom. **(C-D)** UMAP visualization of single-cell RNA sequencing data from E13.5 (C) and E17.5 (D) mouse cortex published in Di Belal et al., 2021. Five major cell clusters, coloured by cell-type assignment based on published metadata (GSE153164), are shown on the left. A combined plot of Unc5D (green) and GPC3 (magenta) mRNA expression per cell is shown on the right. Most of Unc5D-expressing cells belong to the migrating neuron cluster (dashed green line), while GPC3-expressing cells are highly enriched in the apical progenitor (AP) cluster (blue dashed line). Scale bars represent 100 μm (A, B).

**Fig. S6.**
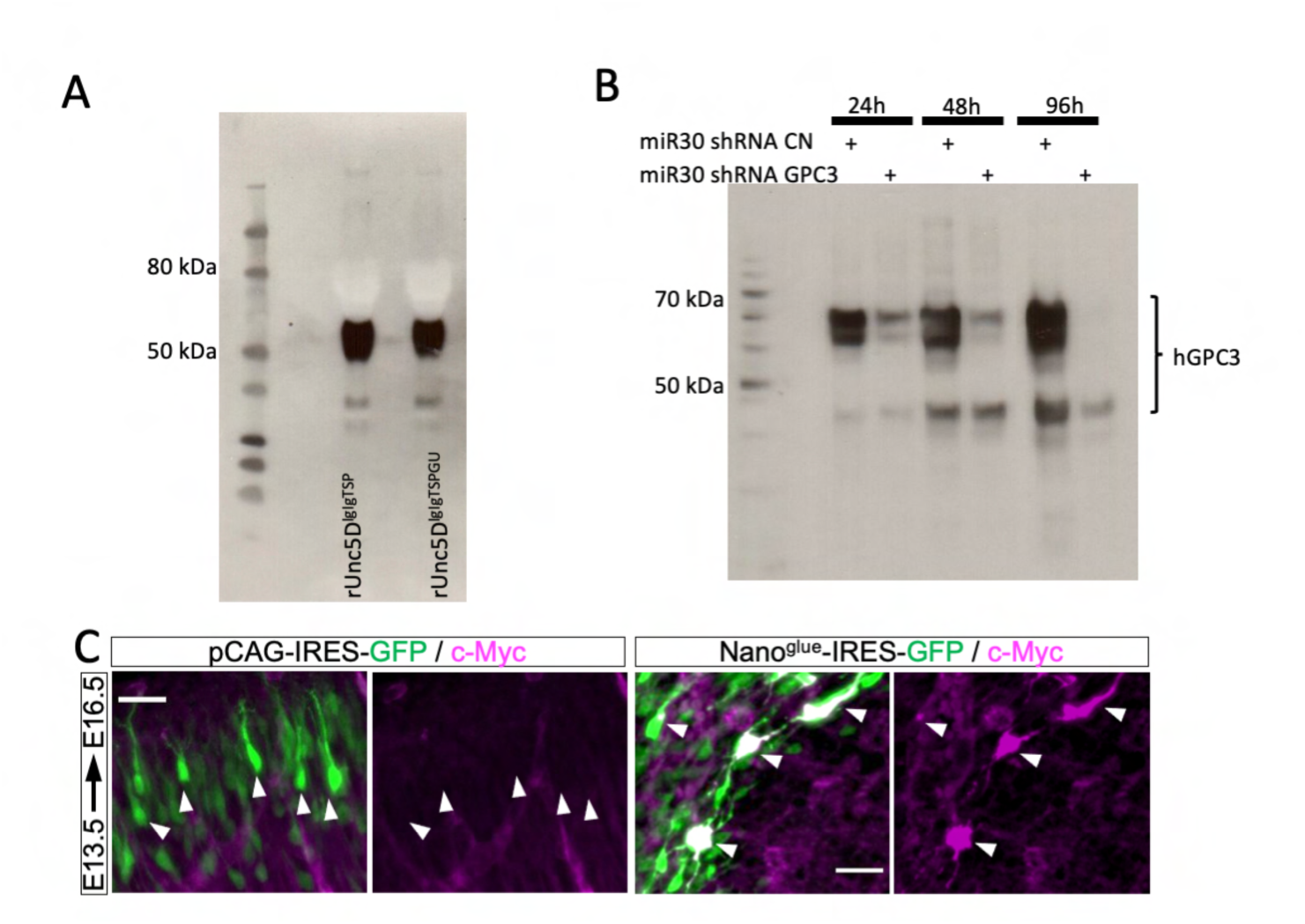
Validation of secreted Unc5D constructs and GPC3 shRNA *in vitro*, nanobody expression *in vivo.* **(A)** Anti-HA western blot showing the secretion levels of HA-tagged rUnc5^IgIgTSP^ constructs that we used in IUE experiments. Supernatants of transfected HEK293 cells were analysed. We find that both constructs are secreted effectively. **(B)** Anti-FLAG blot showing hGPC3 expression in HEK cells, at different time points after transfection (24h, 48h, 96h). HEK cells were co-transfected with vector expressing control (CN) or GPC3 shRNA. Significant reduction in GPC3 expression was observed after 24h, 48h and 96h for cells co-transfected cells with GPC3 shRNA. Similar results were obtained for mGPC3^ecto^ (not shown), as expected, given that the target sequence is conserved in murine and human GPC3. **(C)** IUE of pCAG-IRES-GFP (pCAGIG, control) and pCAGIG encoding Nano^glue^-IRES-GFP was performed at E13.5 and analyzed at E16.5. Myc-tagged Nano^glue^ protein expression in neurons was confirmed by immunostaining with anti-Myc (magenta). Nano^glue^ expression coincides with the positions of cells expressing the reporter GFP (green). White arrows indicate neurons expressing GFP (control and Nano^glue^ plasmid). Scale bar represents 25μm.

**Fig. S7.**
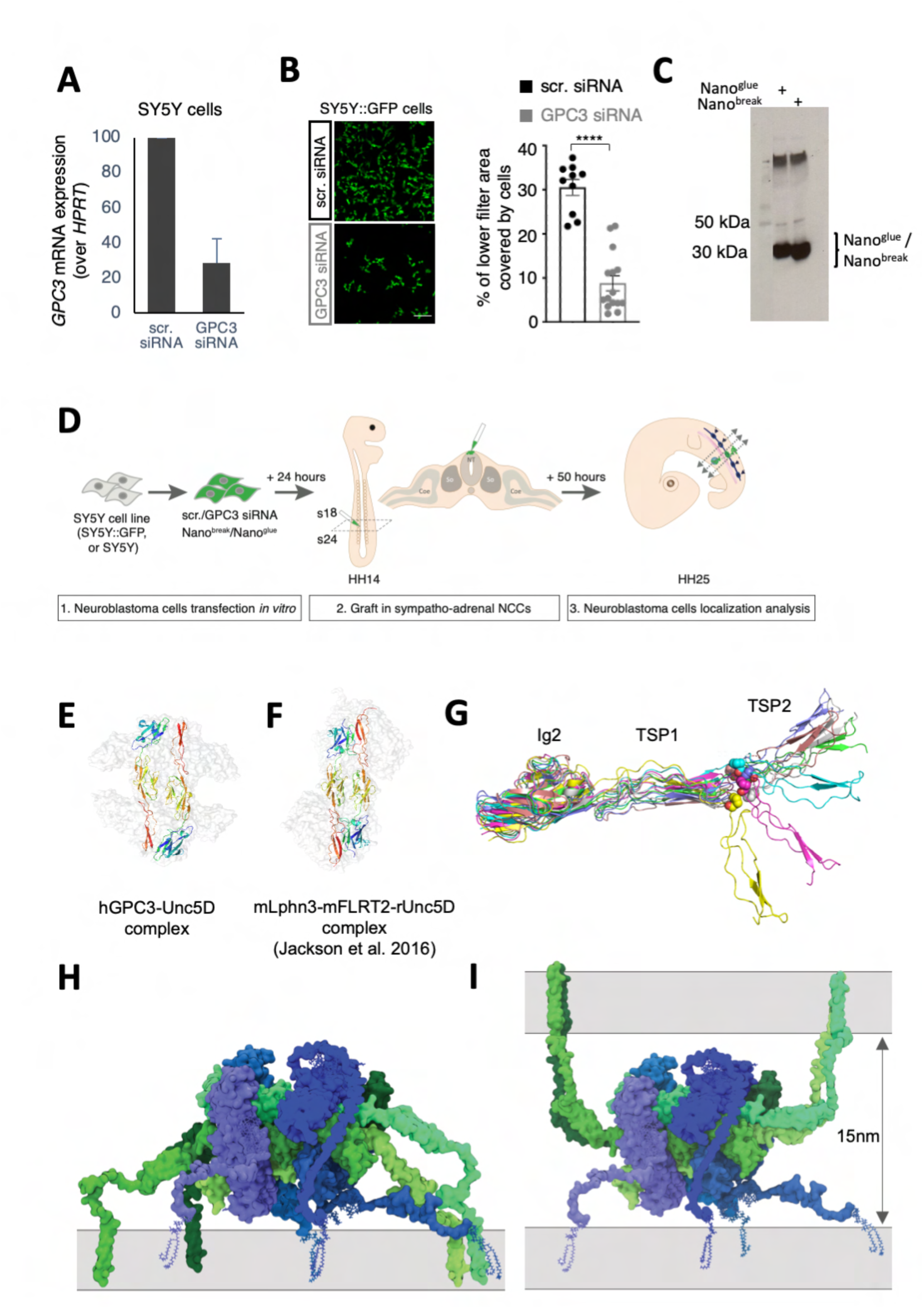
Interfering with GPC3-Unc5 interaction impacts on neuroblastoma cell migration properties. Structural discussion of Unc5 complexes. (A) Q-RT-PCR analysis of GPC3 mRNA expression in SY5Y cells, 24 hours after transfection, using GPC3 siRNA or scr. control. (B) Representative images (left) and quantification (right) of transwell assays measuring the migratory properties of SY5Y:GFP cells transfected with either scr or GPC3 siRNA. ****:p<0.0001. Student T test with Welsch correction. (C) Anti-Myc western blot showing the secretion levels of Myc-tagged nanobody constructs used in Fig. 7F. Supernatants of transfected SY5Y cells were analysed. We find that both constructs are secreted effectively. (D) Scheme of the *in ovo* graft experimental paradigm describing the experiments presented in Fig. 7E, F. (E) Two of the four rUnc5D^IgIgTSP^ chains in the complex with hGPC3^core^ are shown as ribbons, coloured according to the rainbow (N-terminus = blue, C-terminus = red). The rest of the complex is shown as transparent surface (grey). (F) rUnc5D^IgIgTSP^ in complex with FLRT2 and Latrophilin3 (Jackson et al. *Nature Commun.* 2016). The two Unc5D chains are highlighted as rainbow ribbons. (G) Superpositions of Alphafold models of the rUnc5D Ig2-TSP1-TSP2 region, after MD simulation, suggests flexibility in the TSP1-TSP2 linker. (H) The ‘*in cis’* model of hGPC3-rUnc5D was created using Alphafold, MD simulation and MODELLER. GPC3: shades of blue, Unc5D: shades of green. We have not included intracellular domains. (I) As panel H, but showing a potential ‘*in trans’* configuration where GPC3 and Unc5D are expressed on adjacent cells.

**Table S1:**
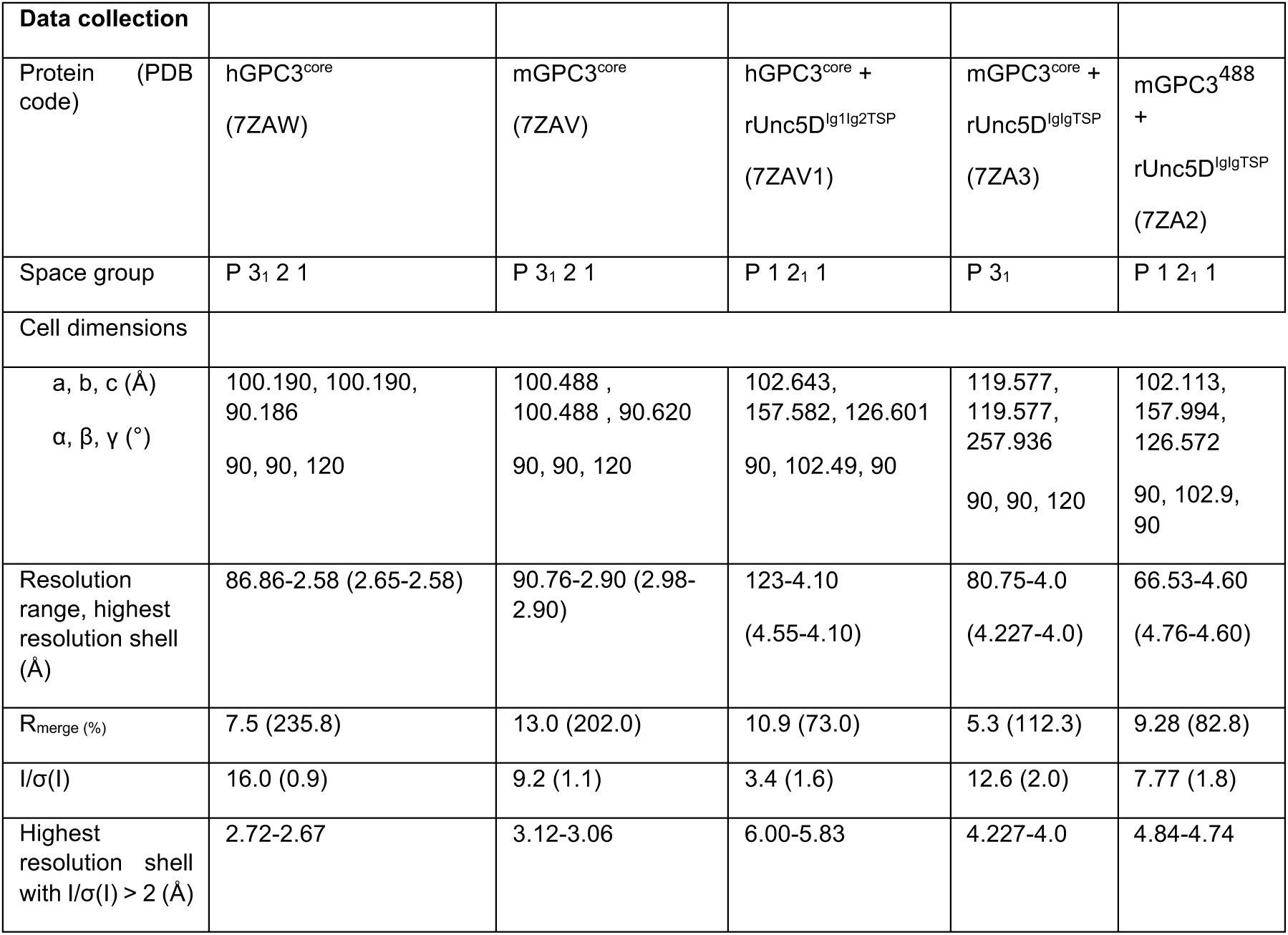
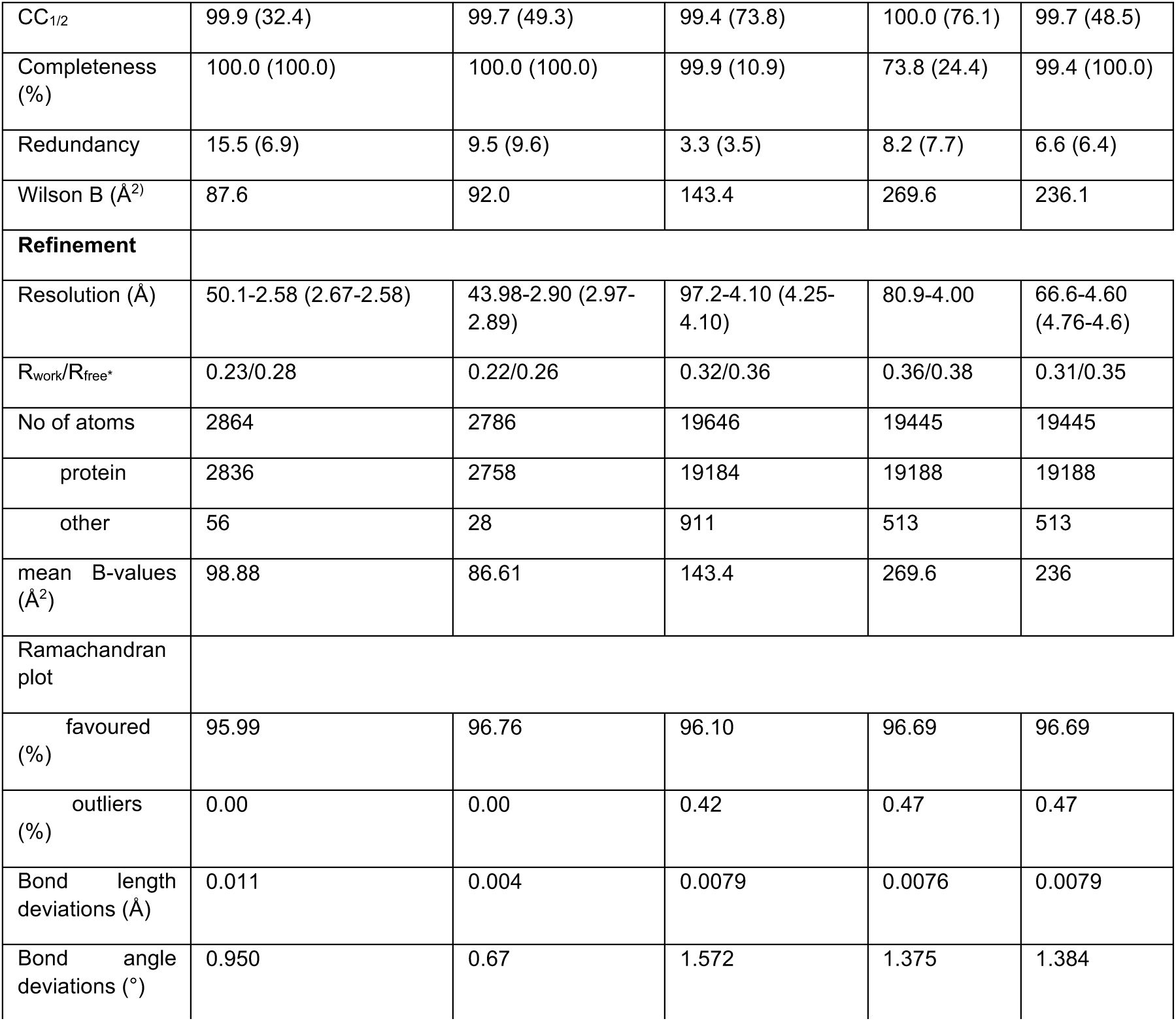
Crystallographic statistics.

